# Identification of chromatin-associated RNAs at human centromeres

**DOI:** 10.1101/2025.06.05.658139

**Authors:** Kelsey Fryer, Charles Limouse, Aaron F. Straight

## Abstract

Centromeres are a specialized chromatin domain that are required for the assembly of the mitotic kinetochore and the accurate segregation of chromosomes. Non-coding RNAs play essential roles in regulating genome organization including at the unique chromatin environment present at human centromeres. We performed Chromatin-Associated RNA sequencing (ChAR-seq) in three different human cell lines to identify and map RNAs associated with centromeric chromatin. Centromere enriched RNAs display distinct contact behaviors across repeat arrays and generally belong to three categories: centromere encoded, nucleolar localized, and highly abundant, broad-binding RNAs. Most centromere encoded RNAs remain locally associated with their transcription locus with the exception of a subset of human satellite RNAs. This work provides a comprehensive identification of centromere bound RNAs that may regulate the organization and activity of the centromere.

## Introduction

Faithful chromosome segregation during cell division is essential for maintaining the genome. In eukaryotes, chromosome segregation is accomplished by attachment of each chromosome to the mitotic spindle through a multiprotein complex, called the kinetochore. The kinetochore is assembled on a specialized chromatin domain, known as the centromere. Kinetochore proteins are conserved across species, however the DNA of the centromere is rapidly evolving. The chromatin of the centromere consists of a core domain that is epigenetically defined by the presence of a centromere specific histone H3 variant, CEntromere Protein A (CENP-A).^1^ The chromatin of the regions surrounding the centromere, known as pericentromeric heterochromatin, is characterized by high levels of the repressive histone modification, H3K9me3.^2^

Human centromeric DNA contains AT rich, tandemly repeated ~171bp monomers of DNA sequence collectively known as alpha satellite DNA.^3,4^ Arrays of tandem repeats within the core centromere are further repeated to form higher order repeats (HORs). CENP-A chromatin occupies a subset of the HORs to generate the active centromere where kinetochores will assemble in mitosis. Recent work has shown that in addition to CENP-A occupancy, the active HOR is also reduced in cytosine DNA methylation compared to the rest of the centromeric DNA.^5^ Satellite repeat arrays flanking HORs, known as pericentromeres, are made up of several different repeat array types, including the major sequence component of classical human satellite fractions I, II, III (HSAT1, HSAT2, HSAT3) originally identified using density centrifugation.^6–9^ Fully sequenced and assembled centromeres from the Telomere-to-Telomere consortium have revealed the complete set of repeat array types that comprise centromeric and pericentromeric regions.^10^ We refer to core centromeric repeats as HORs and the CENP-A occupied region as the active HOR. Pericentromeric repeats denote all other repeat arrays outside of HORs.

In addition to CENP-A, DNA methylation, and histone modification, non-coding RNAs have also been proposed to regulate centromere function. Non-coding RNAs are known to regulate chromatin through several mechanisms. Direct interaction of non-coding RNA with DNA can occur when nascent transcripts hybridize with the template DNA strand forming R-loops or when RNAs form triplex structures with double stranded DNA. R-loops and triplexes have been shown to antagonize or promote DNA methylation by Dnmt3b1 respectively.^11,12^ non-coding RNA also plays important roles in modulating histone modification to repress chromatin through direct regulation of histone methyltransferase complexes, including the HUSH complex,^13,14^ polycomb repressive complex 2 (PRC2),^15–18^ and SUV39H1.^19–21^ RNA has also been implicated in long range genomic organization through interactions with chromatin proteins, including HP1α^22,23^, YY1,^24–26^ and CTCF.^27–31^ However, our understanding of RNA binding by chromatin associated proteins is confounded by potential nonspecific interactions captured by crosslinking dependent immunoprecipitation techniques as well as the broad effects of global RNA degradation leading to debate in the field over whether these interactions are direct RNA binding.^32–34^ Regardless of the interaction mechanism between RNA and chromatin modifiers, it is clear that RNA plays a critical role in genome regulation and organization, particularly with respect to heterochromatin formation.

RNA has also been implicated in chromatin maintenance at both pericentromeric heterochromatin and the core centromere. In mice and humans, RNase treatment resulted in decreased pericentromeric H3K9me3, heterochromatin protein 1 (HP1), and loss of centromeric localization of the histone methyltransferase responsible for depositing H3K9me3, SUV39H1^19,35^ as well as mislocalization of centromeric proteins.^36^ Moreover, SUV39H1 directly binds RNA, and that RNA binding activity is necessary for its localization at pericentromeres.^37–46^ RNA interaction with CENP-C, a necessary component of CENP-A assembly machinery, promotes DNA binding and centromere localization in maize and humans implicating RNA in core centromeric chromatin maintenance.^36,47,48^

Many studies have suggested that RNAs involved in regulating centromeric chromatin may be derived from centromeric repeats. Centromeric transcription and/or centromere derived RNAs have been observed in several species.^49–59^ However, the functional role of centromeric transcription and/or centromeric RNAs is widely debated and differs across organisms. In fission yeast, disruption of the RNAi pathway led to accumulation of centromere derived RNAs and compromised heterochromatin formation.^60^ Deletion of a centromere binding protein in budding yeast led to overexpression of centromeric RNA and decreased expression and centromeric localization of CENP-A.^61,62^ In humans, centromeric transcription has been shown to promote Sgo1 localization to maintain sister chromatin cohesion,^63^ but is not required for CENP-A incorporation at the core centromere.^64^ RNAs transcribed from centromeric and pericentromeric repeats have been implicated in maintaining pericentromeric heterochromatin as well as CENP-A chromatin, and remain associated with their centromere of origin.^65–68^ Abundance of centromeric RNAs is variable across cell types and anti-correlated with centromere proximity to the nucleolus.^69^ Together these studies suggest that centromere-associated RNAs, including those derived from centromeric repeats, are an important component of centromeric and pericentromeric chromatin.

Our understanding of centromere-associated RNAs has been limited by both the lack of unbiased methods to comprehensively identify chromatin associated RNAs and the complex repeat structure of human centromeres which poses a challenge for mapping and localizing sequencing reads. To identify and map RNA-chromatin contacts in a genome wide manner, we developed chromatin associated RNA sequencing (ChAR-seq).^70–73^ ChAR-seq does not require *a priori* knowledge of RNA identity and captures genome scale RNA identity and location information. Here we use ChAR-seq in conjunction with an alignment independent classification of ambivalent sequences with k-mers (CASK)^74^ to identify centromere interacting RNAs. The recent generation of complete human reference genomes with fully assembled centromeres provided a clear picture of the complexity and diversity of centromeric repeats.^10^ Leveraging the T2T assembled centromere sequences we provide a detailed, repeat array level characterization of centromere-associated RNAs. To understand how the RNA-DNA contact patterns at centromeres change across cell types or differentiation states we performed ChAR-seq in the human erythroleukemia cell line, K562, and utilized data from our previous work in human H9 embryonic stem cells (ES) and definitive endoderm (DE) cells differentiated from the ES cells.^73^ Together this study provides the first comprehensive picture of RNA-centromere interactions in humans.

## Results

### Identification and classification of centromere-associated and centromere derived RNAs

We characterized RNA-chromatin contacts in K562 cells using our previously described ChAR-seq approach.^70–73^ Briefly, we generated chimeric molecules capturing RNAs in proximity to DNA by ligating a biotinylated bridge oligo to RNA, digesting the genome with DpnII and then ligating the other end of the bridge to the genome. We next converted the RNA component of each of these chimeric molecules into cDNA. We recovered the bridge containing molecules by precipitation with magnetic streptavidin beads and then amplified and sequenced the molecules **(Figure 1A)**. To map each RNA-DNA contact we split each read into its RNA and DNA derived component and aligned each to the human genome (GRCh38). To annotate repetitive reads, we also performed an alignment independent classification of all RNA and DNA derived sequences using CASK (Classification of ambivalent sequences using k-mers).^74^ CASK generates and intersects k-mer representations of all repeat types to identify k-mers that are unique to each repeat type or are present in multiple repeat types. To differentiate between the different types of centromeric repeat arrays (i.e. Human Satellite 3 vs Higher Order Repeat), we used CASK to generate k-mer databases of T2T assembled centromeric sequences and repeat array type annotations (**Figure 1B**, **Supplemental Figure S1A,B**).^10^ Reads are then classified based on the intersection of all possible repeats in which their k-mers were found. For example, a read that contains k-mers that were only found in a single repeat type, such as HSAT3, can be definitively classified as that repeat type. Reads that contain k-mers that are present in multiple repeat types, and cannot be classified as a single repeat, are annotated with an ambivalence group that encompasses all the k-mer derived repeat types (**Figure 1B**).

**Figure 1:**
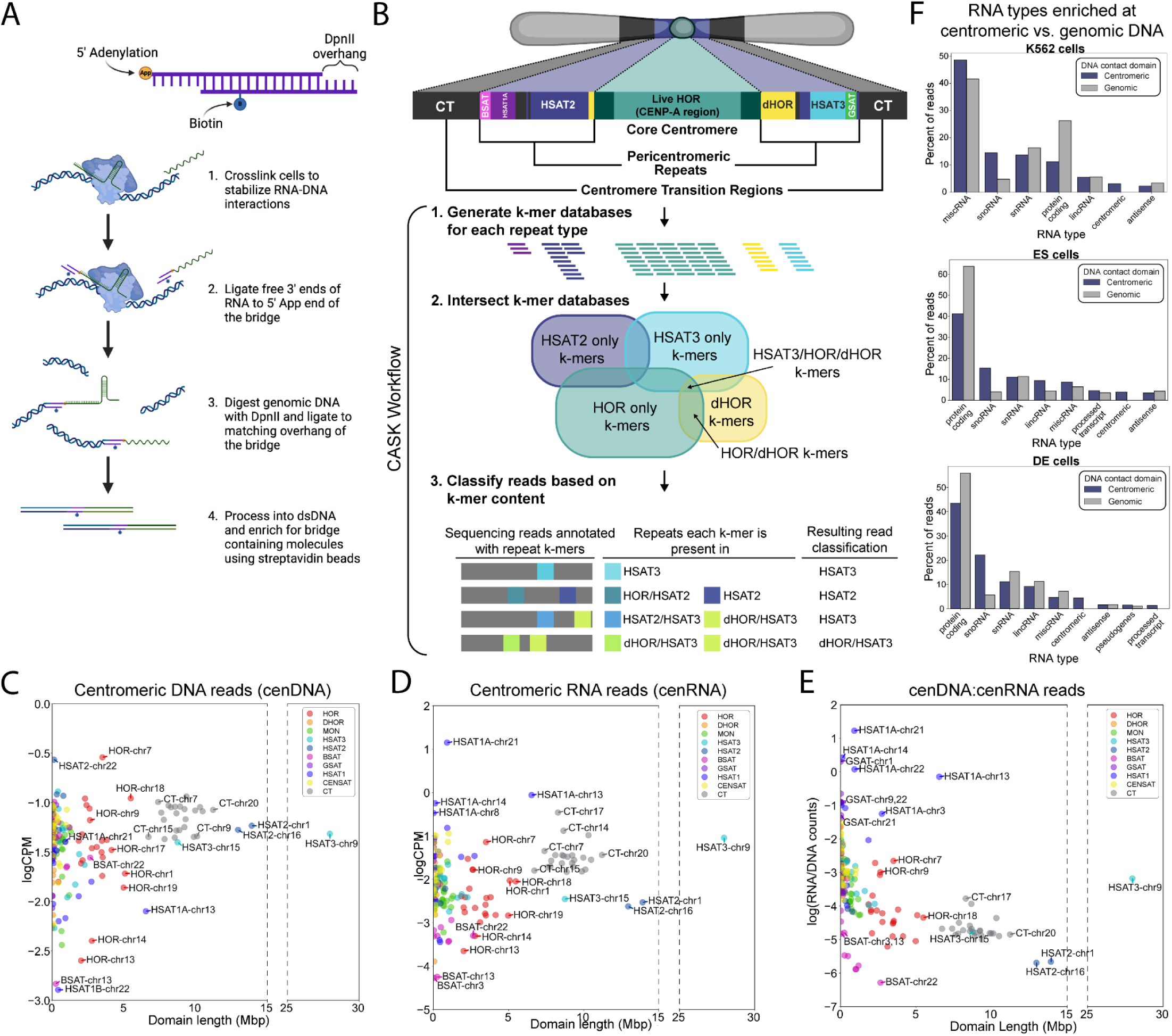
CASK annotation of ChAR-seq identifies centromeric RNA-DNA contacts. **A.** Schematic of ChAR-seq bridge molecule (top) and protocol (bottom). **B.** Schematic of Classification of Ambivalent Sequences (CASK). k-mer databases were generated from T2T assembled centromeric repeat sequences and intersected to create groups of k-mers present in each repeat type or combination of repeat types. Reads were then classified into repeat types of ambivalence groups containing multiple repeat types based on their k-mer content. **C&D.** Log of **C.** DNA or **D.** RNA ChAR-seq reads classified by CASK in K562 cells for each repeat domain normalized by the number of Dpnll sites in the domain per million reads vs. length of repeat domain in Mbp. Reads classified as rDNA regions were excluded. E. Log of ratio of RNA reads to DNA reads classified as each domain normalized by number of Dpnll sites in the domain vs. length of domain in Mbp. Reads classified as rDNA regions were excluded. F. Proportion of RNA reads enriched at centromeric (purple) or non-centromeric (gray) domains belonging to each RNA type in K562 cells (top), ES cells (middle), and DE cells (bottom).

We used CASK to classify both the RNA and DNA derived sequences of ChAR-seq reads in order to examine the contact patterns of 1) RNAs that contact centromeres regardless of their origin and 2) the contact sites across the genome of RNAs that originate from centromeres. To determine how the contact frequency varies across centromeric repeat domains, we counted the number of reads containing centromere derived DNA sequences (cenDNA) regardless of the identity of the RNA derived portion of the read. Accounting for DpnII site frequency and sequencing depth, we found that the number of cenDNA reads classified as each domain varied widely depending on the specific repeat domain and was not correlated with the length of the domain (**Figure 1C**, **Supplemental Figure S2 left panels, Supplemental Table S1**). The same phenomenon was observed for centromere derived RNAs - the number of reads classified as centromeric on the RNA portion of the read (cenRNA) varied across domains and was not correlated with the length of the centromeric domain (**Figure 1D**, **Supplemental Figure S2 right panels, Supplemental Table S1**). For example, the longest repeat array, the human satellite 3 (HSAT3) repeat on chromosome 9, which is 27.9 Mb, contacts roughly the same number of RNAs as the HSAT2 domains that are half the size, and fewer than HORs on chromosome 7 and 18 which are less than one sixth the size. For both cenRNA and cenDNA reads, the relative relationship between most domains remained consistent across all three cell types with the exception of HSAT1A domains. Increased HSAT1A derived RNA reads in K562 cells compared to both ES and DE cells in all replicates may reflect repeat type specific transcriptional regulation (**Figure 1C,D**, **Supplemental Figure S2**). Overall, these data indicate that both the RNA contact frequency at centromeric domains and the contact frequency of centromere derived RNAs vary depending on the identity of the centromeric domain and not its size or cell type.

We next examined the relationship between the number of RNA contacts and the number of RNAs derived from each domain. In previous studies, we observed a correlation between RNA contact density and ATAC-seq peaks which demarcate regions of open chromatin and tend to be more transcriptionally active.^73^ To examine how the relationship between the abundance of RNA contacts and chromatin associated RNAs derived from each domain varied across repeat types, we compared cenDNA and cenRNA counts for each repeat array.

The ratio of cenRNA:cenDNA counts were moderately correlated with the length of the domain (spearman’s rank correlation −0.66) indicating larger domains tend to have fewer RNA reads compared to DNA reads **(Figure 1E, Supplemental Table S1)**. Despite the wide range of cenRNA:cenDNA ratios across centromeric repeat arrays, the relative relationship between arrays remained consistent across replicates and cell types **(Supplemental Figure S3)**. This is most clearly exemplified by HSAT1A arrays which consistently have the highest RNA:DNA count ratios in all cell types (**Figure 1E**, **Supplemental Figure S3**). Together, these data depict features of RNA-DNA contact patterns that are shared amongst members repeat familias (ie. all HORs) that are not entirely dependent on the length of the domain.

### Identification of non-centromere encoded transcripts enriched at centromeric repeats

To identify transcripts encoded by non-centromeric regions that associate with centromeric chromatin, we used CASK to identify ChAR-seq reads in which the genomic DNA was derived from centromeric DNA and genome alignment to annotate the RNA-derived portion of the read. We classified centromeric DNA sequences at the chromosomal level (i.e. all HSAT2 arrays on chromosome 16) and the repeat type level (i.e. the third HSAT2 array on chromosome 16) based on the fully assembled centromeric sequences in the CHM13 T2T genome. The remaining DNA derived portion of the reads that were not classified as centromeric were sorted into 10 kb, 100 kb, and 1000 kb genomic bins. To quantify the enrichment of an RNA (RNA*x*) at a specific genomic region (DNA*x*), we designed an enrichment score (E score) that reflects how frequently RNA*x* contacts DNA*x* relative to its overall contact frequency across the genome. E scores incorporate the number of DpnII sites in the DNA contact domains as well as its length, to normalize for differences in the number of potential contact sites across different DNA contact domains (**Supplemental Figure S4A**). ChAR-seq also captures non-specific RNA interactions with chromatin as any RNA that is diffusing or being transported within the nucleus will make transient contacts with chromatin. To differentiate between bona fide RNA-DNA interactions and nonspecific interactions, we modeled the background contact behavior of RNAs using a generalized additive model.^75^ Because protein-coding RNAs generally do not make specific contacts with the genome outside of their transcription locus, we trained the background model on all protein coding RNA-chromatin contacts, excluding the source chromosome on which the protein coding gene is present. Predicted enrichment scores for each RNA-DNA contact based on the background model were subtracted from observed enrichment scores to generate a residual score (R score). Contacts with residual scores in the top 5% were considered significantly enriched. (**Supplemental Figure S4B,C**).

We identified over 40 non-repeat derived RNAs significantly enriched at centromeres belonging to 8 different classes of RNA. The relative proportion of most classes of RNAs, including snRNAs, miscRNAs, lincRNAs, and antisense RNAs, enriched at centromeres compared to all other genomic regions varied across cell types (**Figure 1F**). However, protein-coding RNAs made up a larger proportion of genome enriched RNAs while snoRNAs consistently made up a larger proportion of RNAs at centromeres (**Figure 1F**). Several snoRNAs and snoRNA host genes, including Gas5 and TMEM107, were enriched at centromeric repeats, but most were enriched at only a subset of centromeric repeat domains, specifically the HSAT2 domains on chromosome 1 and 16. Broad enrichment across multiple centromeric repeat types and chromosomes was only observed for two RNAs - SNORD3A (U3) and RMRP **(Figure 2)**, both of which have been implicated in processing of other non-coding RNAs, particularly in cleavage of pre-ribosomal RNA.^76,77^ RMRP and SNORD3A enrichment at centromeres was observed in all cell types with the strongest enrichment at HORs and HSAT2 domains (**Figure 2**, **Supplemental Figure S5**). Despite the genomic proximity of centromeres on acrocentric chromosomes to rDNA clusters, which are contained within the nucleolus when actively transcribing, centromeric enrichment of nucleolar localized RNAs was not limited to acrocentric chromosomes, but may reflect the close proximity of centromeres to the nucleolar periphery^78,79^ and the potential regulatory crosstalk between the two domains.

**Figure 2:**
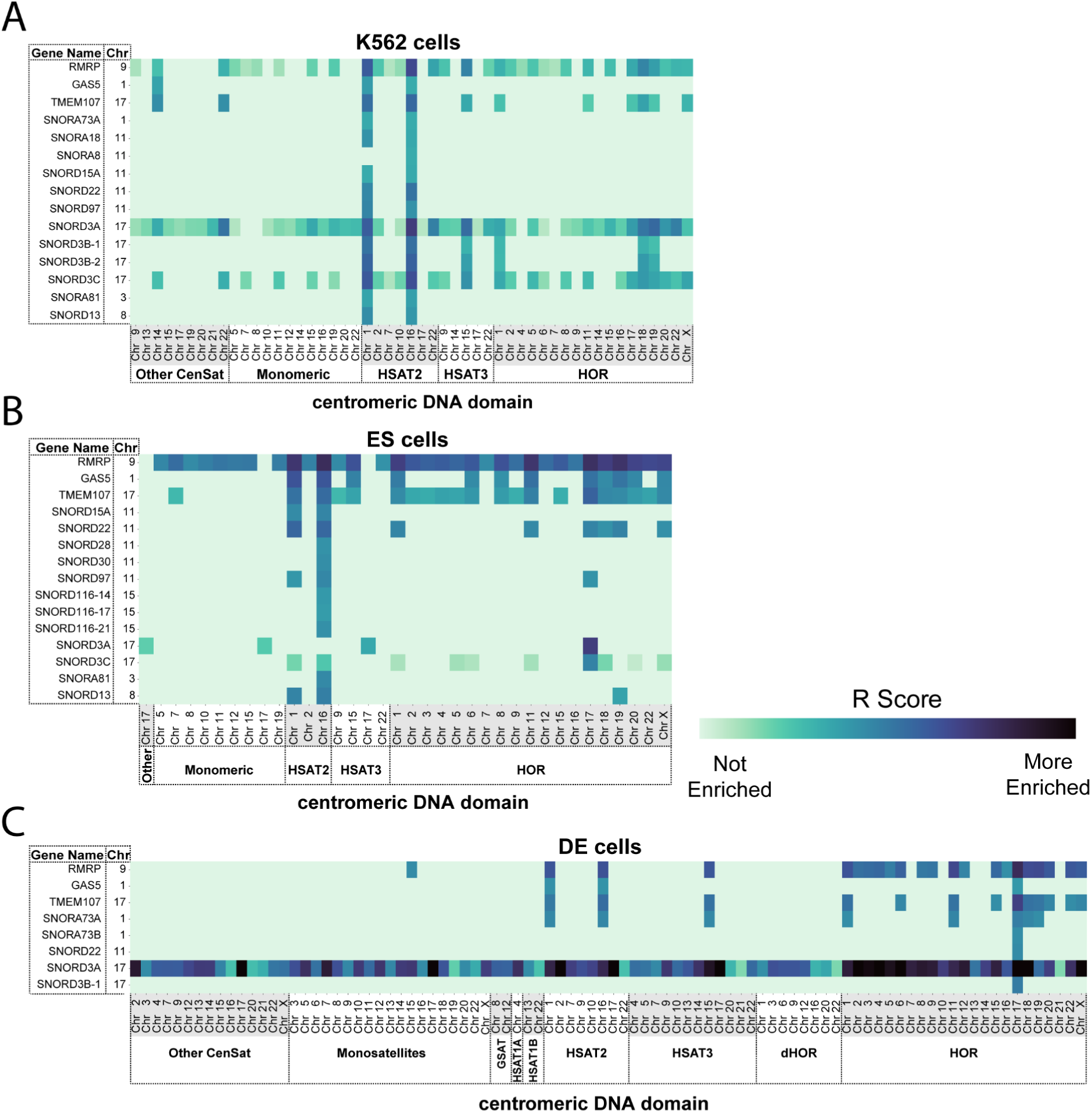
Nucleolar RNAs enriched at centromeric repeats. **A-C.** Heatmap of R scores for nucleolar RNAs (x-axis) enriched at centromeric repeat domains (y axis) in **A.** K562 cells, **B.** DE cells, and **C.** ES cells. DNA side reads were classified into centromeric repeat domains using CASK and are denoted on x-axis with domain type label and chromosome. RNA reads were classified by genomic alignment and denoted with gene name and the chromosome they originate from. Darker color indicates contacts with higher R scores (larger difference between observed number of reads and predicted) indicating stronger enrichment above background contacts. Mitochondrial RNAs, ribosomal RNAs, ribosomal DNA, reads that could not be uniquely assigned to a single repeat type, and all contacts with fewer than 0.2CPM (counts per million total reads) were excluded from heatmap.

Several broad binding RNAs were enriched widely across centromeric and pericentromeric repeat arrays on all chromosomes, including RN7SL1/2 - the RNA component of the signal recognition particle,^80^ several snRNAs involved in regulating splicing, and 7SK - a non-coding RNA that represses RNA polymerase II transcription by binding to the elongation factor P-TEFb **(Figure 3**, **Supplemental Figure S6)**.^80–84^ A subset of previously identified broad binding RNAs, including Y RNAs and Vault RNAs, were also enriched at the centromere, but were limited to HSAT2 domains on chromosome 1 and 16 **(Figure 3**, **Supplemental Figure S6)**. This restricted centromeric enrichment contrasts with their proposed roles in nuclear and cellular processes and widespread nucleoplasmic and cytoplasmic localization. Y RNAs have been shown to regulate RNA stability and DNA replication and have been reported to localize to both the cytoplasm and nucleoplasm as well as directly on chromatin.^85–91^ A small percentage of vault RNAs are associated with vaults - large ribonucleoprotein complexes in the cytoplasm, while the rest are distributed throughout the cytoplasm and are thought to be involved in several processes including nucleocytoplasmic transport and multidrug resistance.^92^ MALAT1, a long non-coding RNA that localizes to nuclear speckles and has been implicated in several processes including transcriptional regulation and splicing,^93^ was enriched widely across centromeric and centromere transition DNA domains only in DE cells **(Figure 3C**, **Supplemental Figure S6C)**. This cell type specific enrichment pattern is consistent with our previous findings that depict MALAT1 as a broad binding RNA that is upregulated in DE cells.^73^ The varied centromeric enrichment patterns of broad binding RNAs illustrate the RNA specific contact behaviors that may be mediated by the inherent properties of the RNA or its interaction partners, as well as by differences in the local chromatin environment across centromeric repeats.

**Figure 3:**
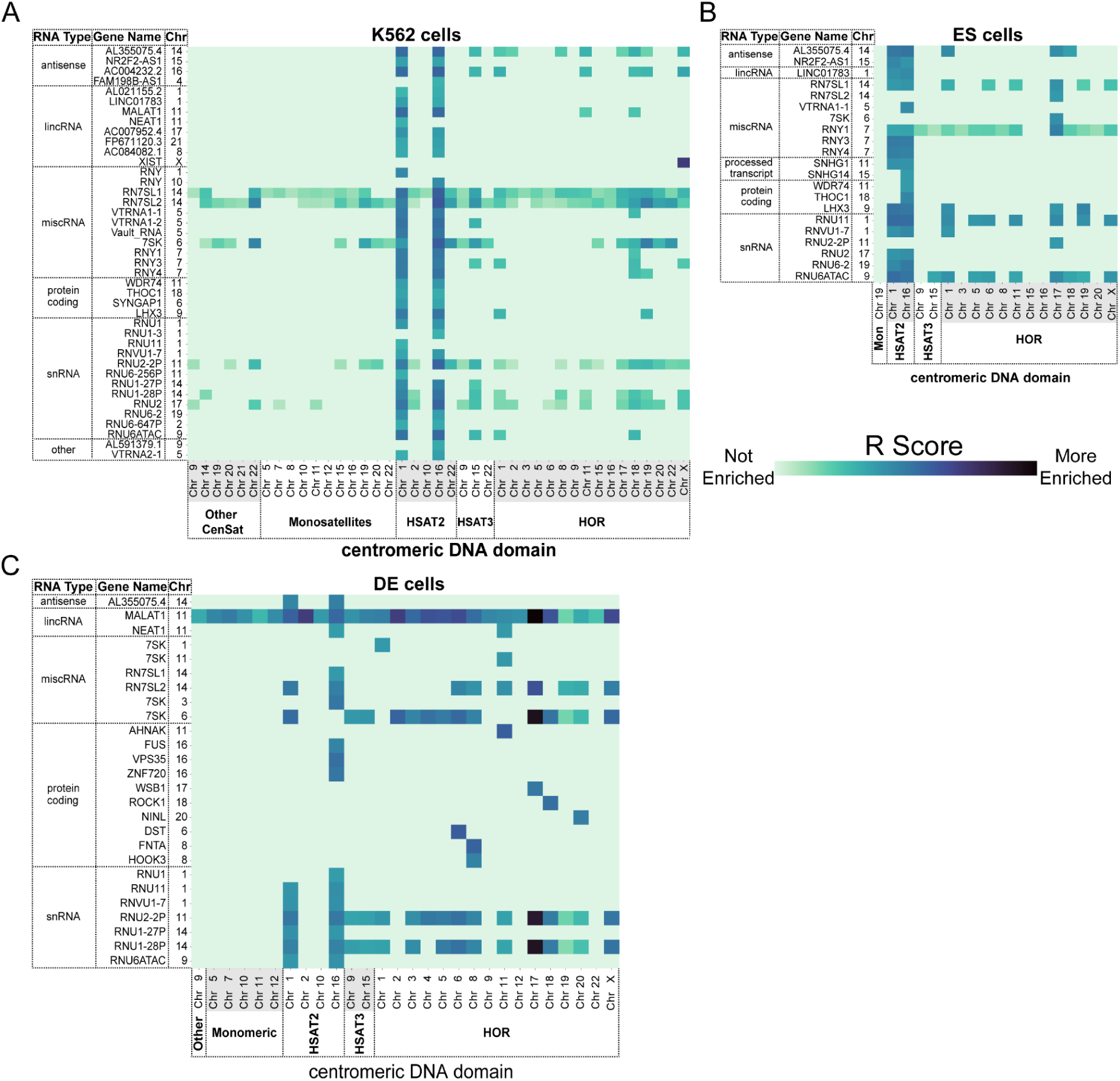
Non-nucleolar localized RNAs enriched at centromeric DNA. **A-C.** Heatmap of R scores for other exon derived RNAs (x-axis) enriched at centromeric repeat domains (y axis) in **A.** K562 cells, **B.** DE cells, and **C.** ES cells. DNA side reads were classified into centromeric repeat domains using CASK and are denoted on x-axis with domain type label and chromosome. RNA reads were classified by genomic alignment and denoted on y-axis with gene type (left most label), gene name (middle label), and the chromosome they originate from (right label). Darker color indicates contacts with higher R scores (larger difference between observed number of reads and predicted) indicating stronger enrichment above background contacts. Mitochondrial RNAs, ribosomal RNAs, ribosomal DNA, reads that could not be uniquely assigned to a single repeat type, and all contacts with fewer than 0.2CPM were excluded from heatmap.

In addition to the RNA specific contact patterns observed at the centromere, individual centromeric repeat arrays display distinct RNA-DNA contact patterns. The most striking repeat array specific patterns are the Human Satellite 2 (HSAT2) arrays on chromosomes 1 and 16 which were found to have many more enriched RNAs than other domains in all three cell lines **(Figure 2,3**, **Supplemental Figure S5,S6)**. Several broad binding and nucleolar localized ncRNAs were only found enriched at these two HSAT2 domains and virtually every RNA enriched at any of the other centromeric repeats was also found enriched at these HSAT2 domains. Given that the HSAT2 domains on chromosome 1 and 16 make up two of the three largest pericentromeric/centromeric arrays (13.2Mbp and 12.7Mbp respectively), we compared the contact behavior of other repeat arrays that spanned over 5Mbp including the longest array, HSAT3 on chromosome 9 which is over 27Mbp and found that no other repeat array of any size displayed similar promiscuous RNA contact behavior (**Figure 2,3**, **Supplemental Figure S5,S6**).

### DNA contact patterns of centromere-derived RNAs

Centromeres are transcribed at low levels in humans but centromeric transcripts play important functions in local regulation of chromatin state. Relatively few centromere encoded RNAs have been studied in depth but in cases where they have been characterized, the RNAs remain locally associated with their sites of transcription.^19,67,68^ To perform an extended analysis of all centromere-associated RNAs we examined the centromere encoded RNAs and their binding sites in the genome. We carried out classification and enrichment calculations of repeat derived RNAs using CASK. We found that most centromere-derived RNAs were exclusively enriched at their transcript locus and did not extend into neighboring domains in K562 and ES cells with increased enrichment at neighboring domains on the same chromosome in DE cells **(Figure 4A-C**, **Supplemental Figure S7,S8)**. Local enrichment of HOR derived transcripts, consistent with previous reports,^67,68^ was observed at the class level which includes all HORs on a given chromosome, as well as the repeat array specific level which describes individual HOR repeat arrays. HOR derived RNAs enriched at the centromere emanated from a single HOR array on each chromosome, typically the active HOR which tends to also be the longest HOR array, and their enrichment generally did not spread even to other HORs on the same chromosome (**Figure 4B**, **Supplemental Figure S7B,D,F**). Overall these data suggest that HOR derived transcripts remain locally associated and do not spread to other centromeres.

**Figure 4:**
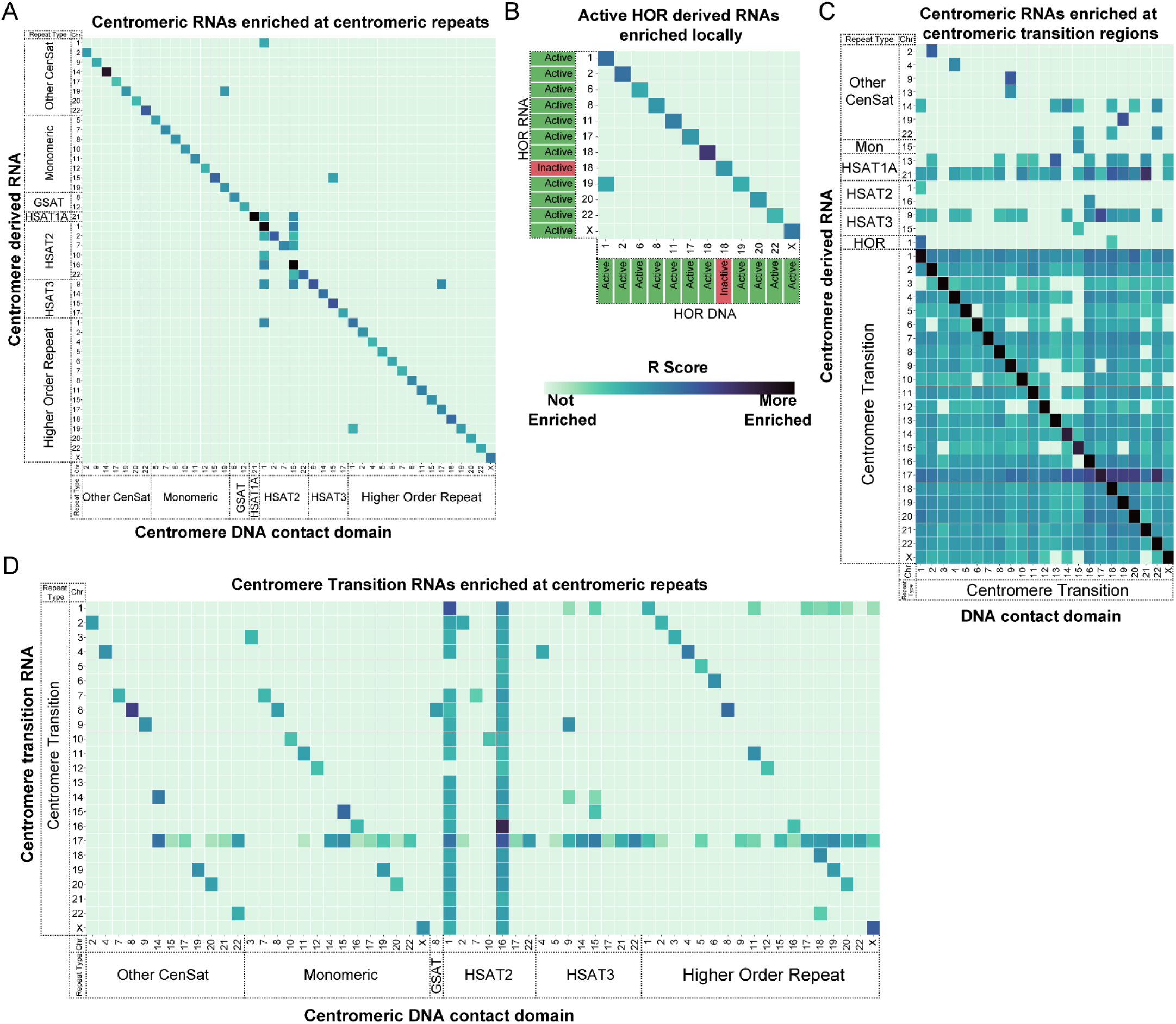
Centromere derived RNAs enriched at centromeric DNA. **A-D.** Heatmap of R scores for RNAs derived from centromere and centromere transition (CT) regions (y-axis) enriched at centromere/CT domains (x-axis) in K562 cells. DNA and RNA reads were classified using CASK and denoted with domain type label and chromosome. Darker color indicates contacts with higher R scores (larger difference between observed number of reads and predicted) indicating stronger enrichment above background contacts. Contacts with R scores below the 95th percentile were considered not significantly enriched and were excluded from heatmaps. Reads classified as ribosomal on either RNA or DNA side, reads that could not be uniquely assigned to a single repeat type, and all contacts with fewer than 0.1CPM were excluded from heatmap. **A.** Class level assigned centromere derived RNAs enriched at centromere DNA domains **B.** Repeat type level HOR derived RNAs enriched at HOR DNA domains. Green label corresponds to T2T active HOR annotation denoting CENP-A containing HOR. Red, inactive label denotes HOR that does not contain CENP-A.C. Centromere/CT derived RNAs (y-axis) enriched at CT regions (x-axis). **D.** CT derived RNAs (y-axis) enriched at centromeric domains (x-axis).

While most centromere derived RNAs displayed a similar local enrichment pattern to that of HOR RNAs, some HSAT and CenSat RNA enrichment did spread to other centromeric domains. The greatest degree of spreading was observed with HSAT3 and HSAT1A derived transcripts which were enriched at HSAT2 domains on chromosome 1 and 16 and across all centromere transition (CT) regions which are genic regions flanking pericentromeric repeats. Centromeric transcripts that were enriched outside their own transcription locus were often also enriched at HSAT2 domains on chromosomes 1 and 16 **(Figure 4A)** which appear to act as RNA contact hubs for centromere derived transcripts in addition to non-repetitive RNAs **(Figure 2,3,4)**. Despite the increased contacts at HSAT2 DNA domains **(Figure 2,3,4A,C)**, the abundance of reads with HSAT2 derived RNA sequences was equal to or less than that of most other domains **(Figure 1D)** and enrichment of RNAs derived from HSAT2 domains on chromosomes 1 and 16 generally did not spread outside HSAT2 domains **(Figure 4A,C**, **Supplemental Figure S7 left panels, S8A-E)**. The distinct behavior of different HSAT repeat types illustrates that the RNA contact behavior of a centromeric domain and RNAs derived from it are features specific to the repeat type and not general features of centromeres or pericentromeres.

RNAs derived from genic centromere transition (CT) regions exhibited a distinct pattern with enrichment across CT regions on every chromosome as well as other centromeric repeat domains on the same chromosome **(Figure 4C,D**, **Supplemental Figure S8,S9)**. RNAs derived from centromere transition regions on chromosome 17 were broadly enriched across centromeric domains **(Figure 4D**, **Supplemental Figure S9)** mimicking the pattern seen with SNORD3A which is encoded within the centromere transition regions on chromosome 17, likely representing the overlap between CASK assignment and genome alignment in non-repetitive regions. This widespread enrichment could be indicative of increased transcriptional activity as well as increased accessibility allowing for more RNA contacts. Additionally, the uniqueness of CT regions results in a larger database of CT specific k-mers increasing the proportion of reads that can be confidently classified as CT **(Supplemental Figure S1A)**.

## Discussion

By analyzing RNA-DNA contacts across all centromeres we provide the first comprehensive description of centromere enriched RNAs in human cells. We used ChAR-seq data that links RNAs to their binding sites in the genome to identify RNAs that bind to centromeres and centromere encoded RNAs that bind across the genome. Applying CASK, which classifies sequences based on their repeat content, to annotate centromeric reads in CHAR-seq data, we demonstrate that RNA-DNA contact patterns differ drastically between repeat array types. Understanding the RNA contact landscape at centromeres expands our definition of the complex and specialized chromatin environments at centromeres beyond what is described by differences in DNA sequence and histone content.

We hypothesized that RNAs necessary for establishing core centromeric chromatin would be enriched at every centromere. We identified a subset of nucleolar localized ncRNAs, including SNORD3A and RMRP, that were enriched at centromeres on virtually every chromosome. SNORD3A (U3) is a highly abundant box C/D snoRNA that, in contrast to most other box C/D snoRNAs, does not mediate RNA methylation, but instead has been shown to base pair with pre-ribosomal RNA to direct site specific cleavage.^94–97^ SNORD3A has also been reported to competitively bind an miRNA responsible for down regulating uridine monophosphate synthetase (UMPS) expression and has also been shown to encode several of its own miRNAs.^98,99^ RMRP, the RNA component of the mitochondrial RNA processing endoribonuclease, has been shown to cleave mitochondrial RNA as well as ribosomal RNA.^77,100^ Mutations in RMRP have been linked to changes in gene expression and defects in cell cycle progression in the disease state cartilage-hair hypoplasia.^101^ This enrichment pattern echoes previous reports of centromere and peri/centromeric RNA localization at the nucleolar periphery.^59,68,78,79^ The bias for snoRNA enrichment at HORs compared to other centromeric repeats may be further evidence of a regulatory role for the interaction between centromeres and the nucleolus.^68^ In addition to their broad centromere enrichment, the proposed multifunctionality of SNORD3A and RMRP make them compelling candidates for centromere regulation. The only other RNAs that were enriched across most centromeres were abundant broad-binding RNAs involved in transcriptional regulation, splicing, and RNA modification. We did not detect CCTT RNA which was previously reported to localize at centromeres and bind CENP-C.^102^ In light of these observations, we propose that centromere enriched nucleolar RNAs, especially SNORD3A and RMRP, represent the most compelling candidates for deeper investigation into their potential roles in chromatin regulation.

The vast majority of enriched RNAs at centromere are found contacting the HSAT2 domains on chromosomes 1 and 16 which seem to act as RNA contact hubs. Each array spans over 12 Mbp making them the largest HSAT2 arrays and among the largest repeat arrays across all centromeres. The other largest arrays only have a fraction of the enriched RNAs indicating that the increased enriched contacts cannot be simply explained by array length. In contrast to their promiscuous RNA contact behavior, RNAs derived from these HSAT2 domains were mostly enriched at their transcription locus with some spreading that mimics the contact pattern of other HSAT and CenSat derived RNAs in that cell type. The RNA-DNA contact pattern at these massive HSAT2 arrays depict a chromatin environment that is unique from all other repeat domains and may be evidence of repeat array specific epigenetic regulation.

Despite differences in relative abundances, HOR derived RNAs are almost exclusively enriched at their own transcription locus. While the function and importance of active transcription at the centromere is debated and differs between organisms, the presence of HOR derived transcripts and their *cis* localization is in agreement with previous work^19,67,68^ and implies that if HOR derived RNAs play a role in centromeric chromatin regulation, it is mediated by RNA *in cis* and not by a single or subset of HOR RNAs diffusing between centromeres. This does not appear to be a centromere wide phenomenon as HSAT1A, HSAT3, and some monomeric satellites (Mon), and other centromeric satellites (CenSat), and especially CT derived RNAs do spread to other repeat arrays both on their own chromosome and to others.

The different RNA contact patterns observed across repeat types in conjunction with the increased spreading of centromere derived RNAs seen in DE cells illustrates the repeat array and cell type specific contact patterns. Locus specific RNA-DNA contact patterns may be a component of the distinct local chromatin environments that exist with the centromere and surrounding pericentromeric regions. Differences in RNA-DNA contacts across cell types may also reflect the changes in chromatin environment that occur through differentiation. ES and many cancer cell lines, including K562, have been shown to have less constitutive heterochromatin, particularly at pericentromeres.^103–107^ Increased heterochromatin in DE cells may impact 3D genome organization either through compaction or proximity to the nucleolus or other RNA hubs and would result in the increase in cis and trans RNA contacts observed at the centromere. Further study in cells with compromised heterochromatin maintenance is necessary to delineate the role of compaction and 3D genome organization in centromeric RNA-DNA contacts.

This study describes an RNA-chromatin environment that is rarely uniform across centromeric repeat types or cell types. It is clear from this work that to advance our understanding of the role of RNA at the centromere, it is necessary to consider each subdomain of the centromere or pericentromere and the unique environment they occupy individually. The comprehensive list of centromere-associated RNAs and the specific domains they are enriched at has provided the groundwork for further investigation into the role of RNA in regulating centromeric chromatin. Future work to selectively perturb these centromere-associated RNAs and/or centromere derived RNAs will provide insight into their functional contribution to centromeres.

## Methods

### Cell harvest and fixation

Human K562 cells were cultured in RPMI 1640 media supplemented with 50 U/mL penicillin and streptomycin to a concentration of 0.8-1×10^6^ cells/mL. For each sample, 10×10^6^ cells were collected in 50 mL conical tubes, centrifuged at 500 xg for 5 min, liquid media was aspirated, and resuspended in 16.25 mL serum free media. Cells were fixed by adding 3.75 mL fresh 16% formaldehyde while shaking to a final concentration of 3% and incubated on a rocker for 10 minutes at room temperature. Formaldehyde was quenched by adding 6.25 mL of 2.5 M glycine and incubating on rocker at room temperature for 5 minutes and on ice for 15 minutes with intermittent mixing. Fixed cells were then pelleted and resuspended in ~1 mL ice cold PBS. Using a hemocytometer, cells were counted and aliquoted to 10 million cells each.

### ChAR-seq library preparation

Two replicate ChAR-seq libraries were generated for K562 cells as in(Limouse et al 2023) with the following modifications: 1) After decrosslinking, DNA was isolated by phenol:chloroform extraction. Briefly, 300 μL phenol:chloroform:isoamylalcohol was added to each sample and vigorously mixed. The entire volume was transferred to a heavy phase lock tube and centrifuged at 13,000 x g for 3 minutes at room temperature. 300 μL chloroform was added to the top layer while still in the phase lock tube, samples were inverted several times to mix and then centrifuged again at 13,000 x g for 3 minutes at room temperature. The top layer was transferred to a fresh 1.5 mL microcentrifuge tube with 900 μL 100% ice cold ethanol, incubated at −20°C for 20 minutes, and centrifuged at max speed for 20 minutes at 4°C. Ethanol was aspirated, and DNA pellet was washed with 80% ethanol, spun again at max speed for 5 minutes at 4°C. All ethanol was removed and samples were allowed to air dry before the DNA pellet was resuspended in 132 μL TE (10 mM Tris-HCl pH 8, 1 mM EDTA pH 8). 2) Two additional wash steps were performed after initial binding to magnetic streptavidin beads (for a total of 4 washes), one at 50°C for 2 min, and one at room temperature. 3) The optional PacI digest was performed as described. 4) Libraries were quantified using high sensitivity Qubit Fluorometer 3.0 and size distribution was determined using high sensitivity Agilent TapeStation 4200. Libraries still containing high or low molecular weight fragments underwent an additional size selection using SPRIselect paramagnetic beads (BeckmanCoulter B23317) prior to sample pooling. 5) Libraries were first sequenced at a low depth using an illumina MiSeq to assess bridge content and duplication rate prior to pooling and high depth sequencing using illumina NovaSeq X Plus. After sequencing, raw data files for each replicate were combined. All downstream processing was performed on combined files.

### ChAR-seq data processing

Raw reads were processed using a snakemake pipeline described in Limouse et al. 2023.^72^ Briefly, PCR duplicates were removed, sequencing adapters were trimmed, and reads lacking any bridge sequence were filtered out. The remaining reads were split into RNA and DNA fastq files corresponding to the sequences of the RNA-derived and DNA-derived portions of the chimeric molecules and aligned to hg38 transcriptome and genome respectively. RNA reads aligned to ribosomal sequences and corresponding DNA sides were filtered out.

### Repeat classification

CASK (classification of ambivalent sequences using k-mers). First, repeat sequence k-mer databases were generated using KMC (https://github.com/refresh-bio/KMC) for each repeat class and repeat type based on repeat sequences extracted from genome fasta file (T2T CHM13v2; https://github.com/marbl/CHM13) defined by genomic coordinates (Cen/Sat bedfile from T2T CHM13v2) using bedtools. A reference repeat k-mer database was then generated by aggregating all repeat k-mers, subtracting k-mers present outside of defined repeat regions, and assigning each k-mer a unique identifier using BBDuk (part of BBTools suite). Reference repeat k-mers were then identified and annotated in both RNA and DNA ChAR-seq reads using BBDuk. Reads were then classified into ambivalence groups based on the intersection of all possible repeats defined by k-mer content. For example, a read containing 3 different repeat k-mers, where the first is present in repeat_type1 and repeat_type2, the second is present in repeat_type1 and repeat_type3, and the third is present in repat_type1 and repeat_type4, would be classified as repeat_type1. Reads were then counted for each RNA-DNA pair. Non-repetitive RNA reads were annotated according to their alignment to hg38 and non-repetitive DNA reads were annotated with a genomic bin identifier. Normalized CPM (counts per million reads) were calculated as the number of RNA or DNA reads annotated as each repeat class (N), normalized by the number of DpnII (D) sites present in the repeat domain as well as by the total number of reads in the dataset after filtering (R) per million reads (R/10^6^): (N/D)/R/10^6^.

### Enrichment calculation

To quantify RNA contacts at repetitive regions, we calculated an enrichment (E score) for each RNA-DNA contact as the number of times that RNA-DNA combination was observed in the dataset normalized by the number of DpnII restriction sites in the DNA contact domain. To account for the fact that more highly expressed RNAs and larger DNA domains will make more contacts, these counts were also normalized by the total number of times that RNA was found contacting anywhere in the genome, and the length of the contact domain **(Supplemental Figure S4A)**.

The background contact behavior of RNAs non specifically interacting with the genome was established using a generalized additive model of all trans contacting protein coding RNAs. For each RNA-DNA contact, a predicted enrichment score was calculated based on the background model specific for the contact chromosome which includes all protein coding RNAs contacting that chromosome and transcribed from any other chromosome (trans contacting). For DNA contacts including ambivalence groups, the background model included all trans contacting protein coding RNAs for all chromosomes in the ambivalence group. Predicted enrichment scores for each RNA-DNA were then subtracted from observed enrichment scores to generate a residual score (R score). The top 5% of residual scores were considered significantly enriched and plotted in heat maps **(Supplemental Figure S4B,C)**.

## Data and software availability

ChAR-seq data previously generated in embryonic stem cells and definitive endoderm cells are available as GSE240435. K562 ChAR-seq data generated as a part of this study are available through SRA as PRJNA1270001. Processed data supporting the findings of this study for all cell types are available as GSE298896.

All software packages and code released as part of this study are available as public repositories https://github.com/straightlab/. Software packages as well as Snakemake pipelines used to process raw ChAR-seq data are available at https://github.com/straightlab/chartools and https://github.com/straightlab/charseq_dynamics_paper. CASK repeat classification tool is available at https://github.com/straightlab/cask. Enrichment calculation pipeline and all code used to generate figures in this study are available at https://github.com/straightlab/centrochar.

## Acknowledgements

We thank the members of the Straight lab for meaningful discussion and feedback, and Pragya Sidhwani for comments on data analysis and figures. The computing for this project was performed on the Stanford Sherlock HPC cluster and we thank Stanford University and the Stanford Research Computing Center for providing computational resources and support that contributed to these research results.The development of the ChAR-seq method and CASK tools was supported by NIH/NHGRI R01HG009909 to AFS. This work was supported by NIH R01GM074728 to AFS and NIH Training Grant T32-GM007790-40 to KAF.

## Author Contributions

K.A.F: generated ChAR-seq libraries for K562 cells, conducted data analysis, prepared this manuscript, and generated figures. C.L. designed and wrote ChAR-seq data processing pipeline and k-mer classification software. A.F.S. supervised the work and contributed to manuscript preparation and editing.

## Declarations of interest

The authors declare no competing interests.

## Declaration of generative AI and AI-assisted technology

During the preparation of this manuscript, the authors used ChatGPT to suggest alternative words and phrases in some sentences to improve clarity. The authors reviewed and edited all content and take full responsibility for the content of the publication.

## Supplemental Information

Document S1: Supplemental Figures S1-S9

Table S1: excel file containing all correlation statistics for CPM comparisons.

---

**Supplemental Figure S1:**
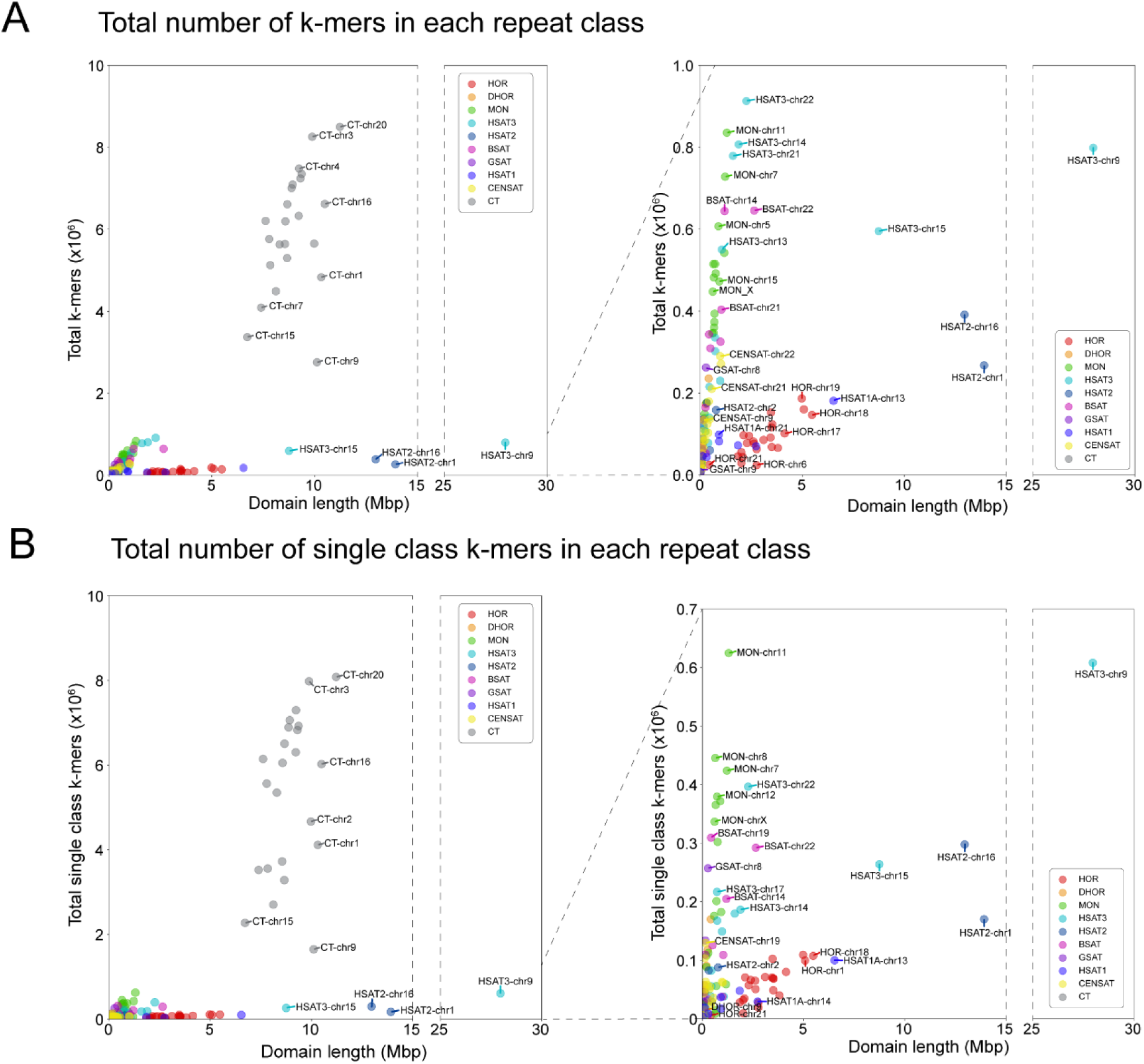
**A.** Scatter plot depicting the total number of different k-mers in each repeat class (y-axis) vs. class length in Mbp. Includes k-mers that are shared between multiple repeat classes as well as k-mers that are class specific. Right plot shows zoomed in view. **B.** Scatter plot depicting the total number of different k-mers unique to each repeat class (y-axis) vs. class length in Mbp. Includes k-mers unique to single repeat arrays (i.e. HOR-chr1-1: the first HOR domain on chr1) and unique to classes (i.e. HOR-chr1). Right plot shows zoomed in view.

**Supplemental Figure S2:**
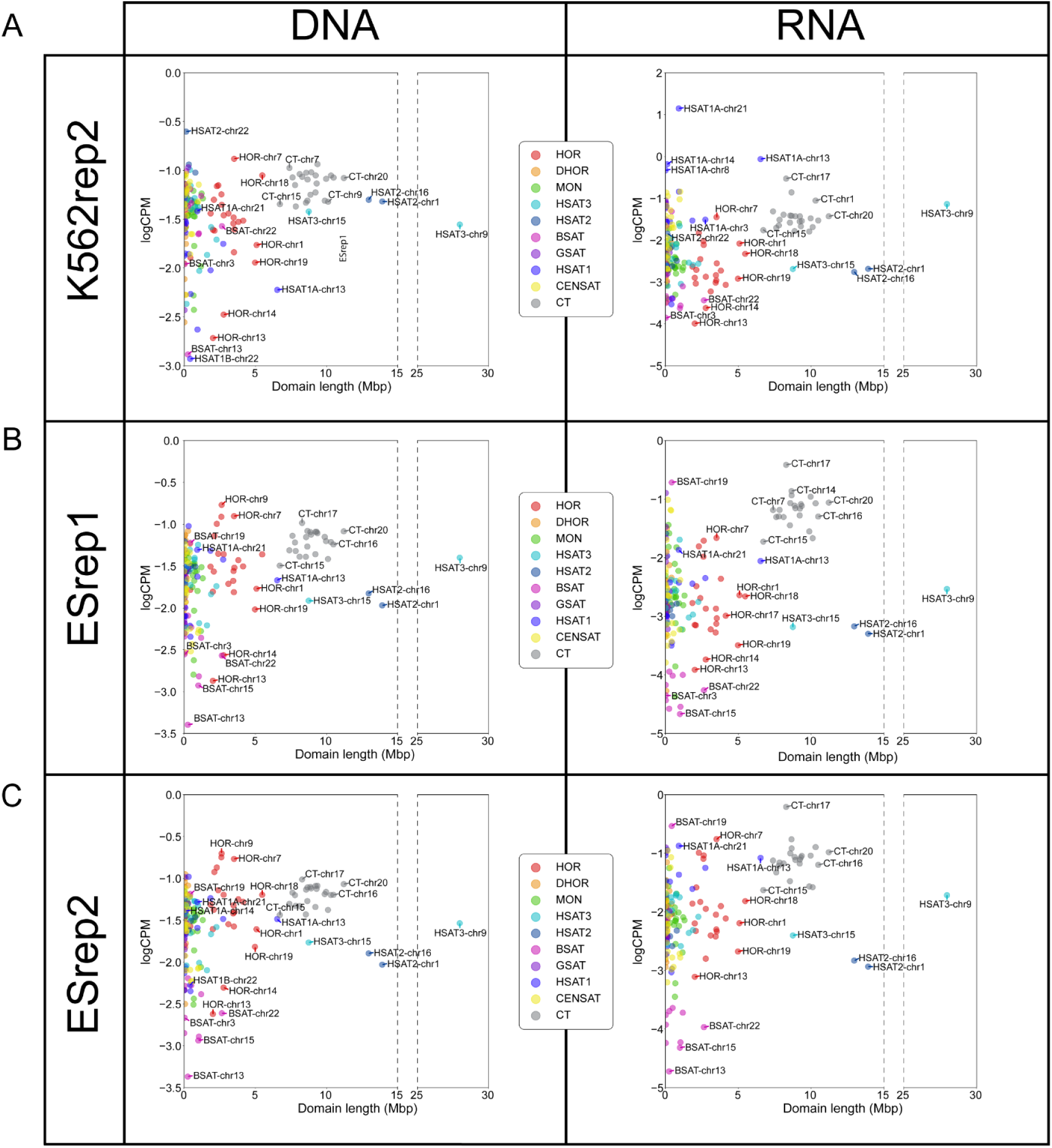
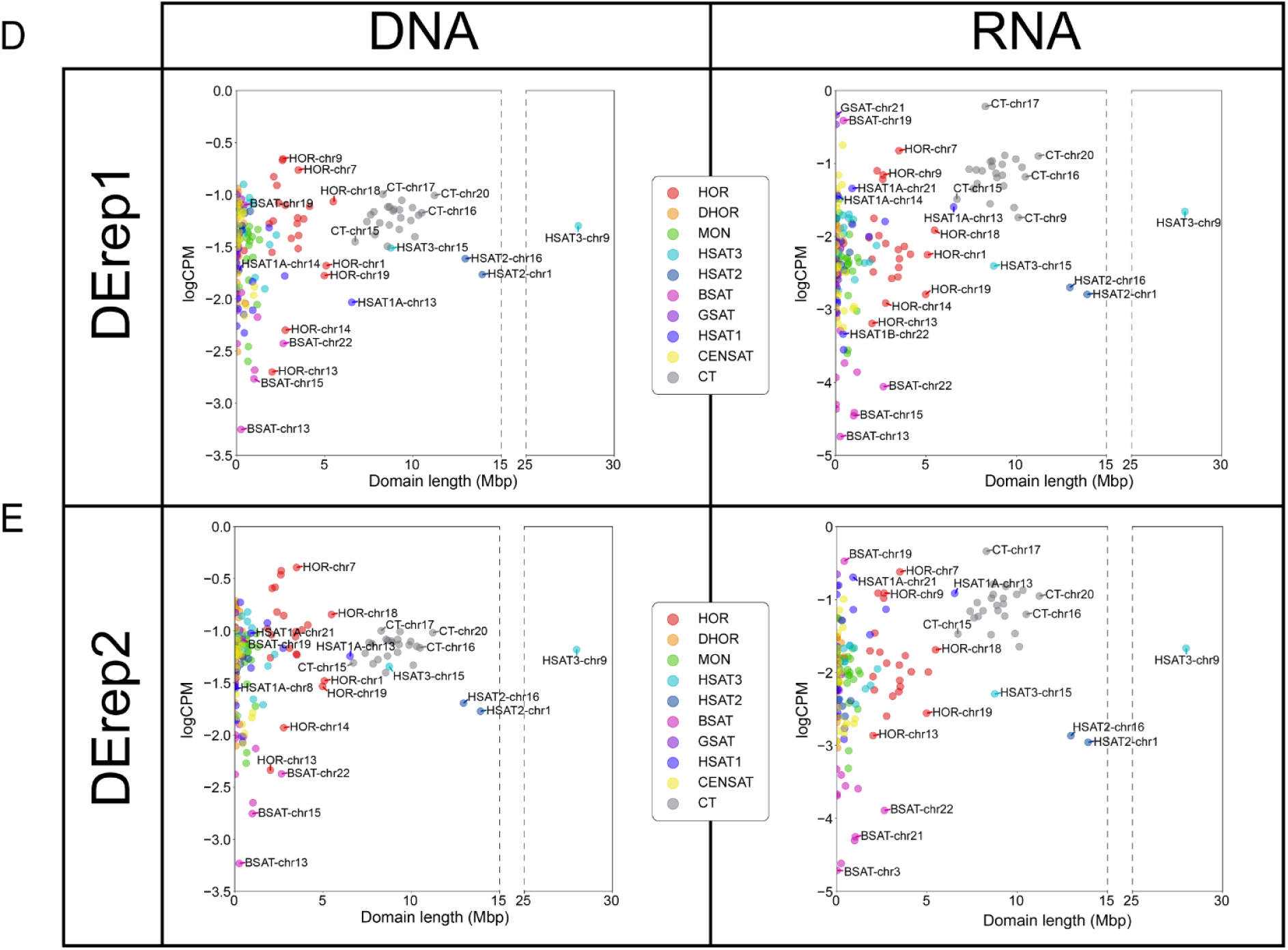
Normalized cenDNA and cenRNA read counts vs. domain length: **A-E.** Similar to Figure 4.1C&D: Log of DpnII normalized DNA and RNA read counts classified by CASK for each repeat domain per million reads. Reads that could not be uniquely classified or were classified as rDNA were excluded. **A.** K562 replicate 2 DNA reads (left) and RNA reads (right) **B.** ES replicate 1 DNA reads (left) and RNA reads (right) **C.** ES replicate 2 DNA reads (left) and RNA reads (right) **D.** DE replicate 1 DNA reads (left) and RNA reads (right) **E.** DE replicate 2 DNA reads (left) and RNA reads (right)

**Supplemental Figure S3:**
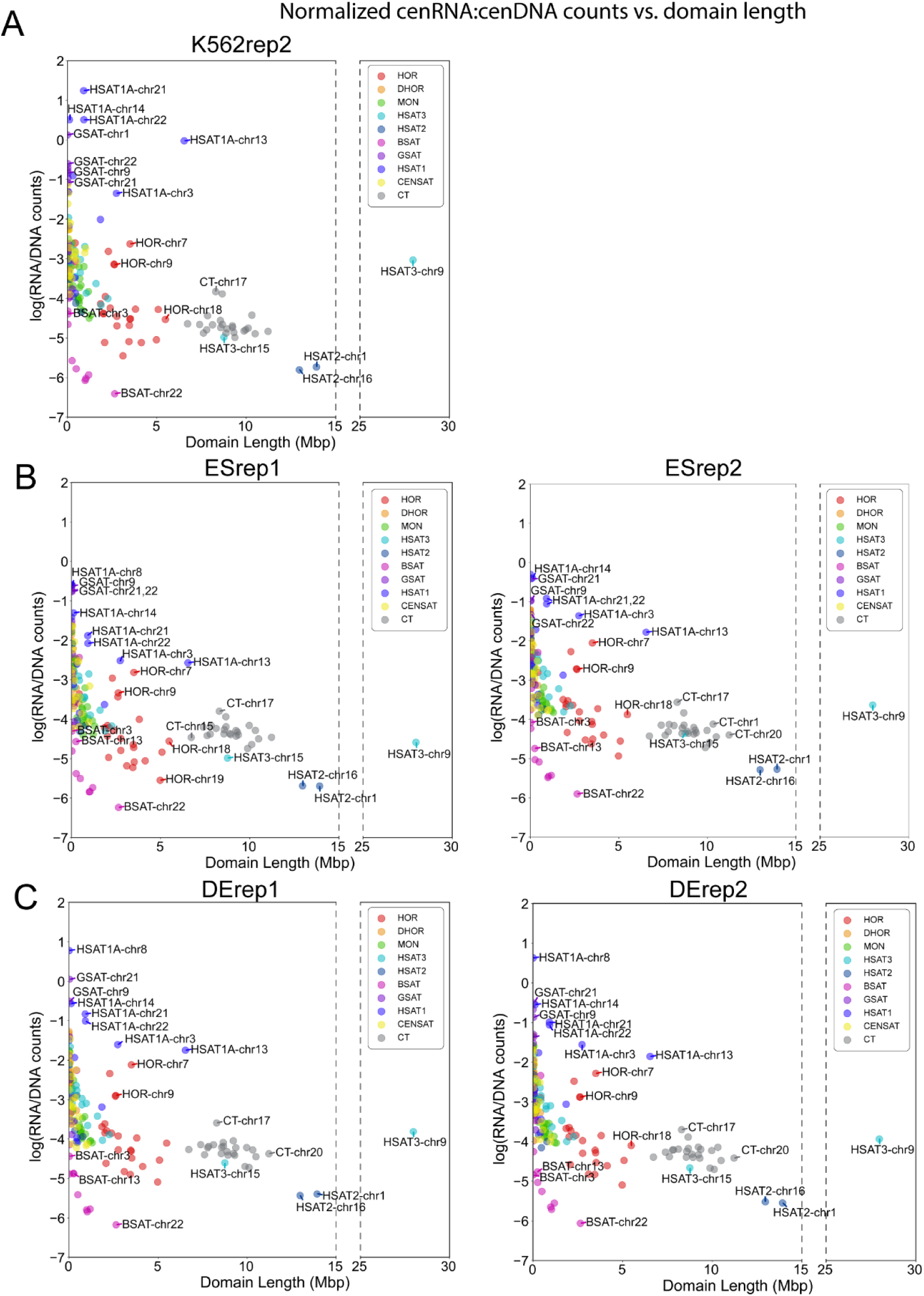
Normalized cenRNA:cenDNA counts vs. domain length. **A-C.** Log of ratio of RNA to DNA read counts classified by CASK for each repeat domain divided by the number of DpnII sites in the domain. Reads that could not be uniquely classified or were classified as rDNA were excluded. **A.** log(RNA:DNA) for K562 replicate 2 **B.** log(RNA:DNA) for ES replicate 1 (left) and ES replicate 2 (right) **C.** log(RNA:DNA) for DE replicate1 (left) and DE replicate 2 (right)

**Supplemental Figure S4:**
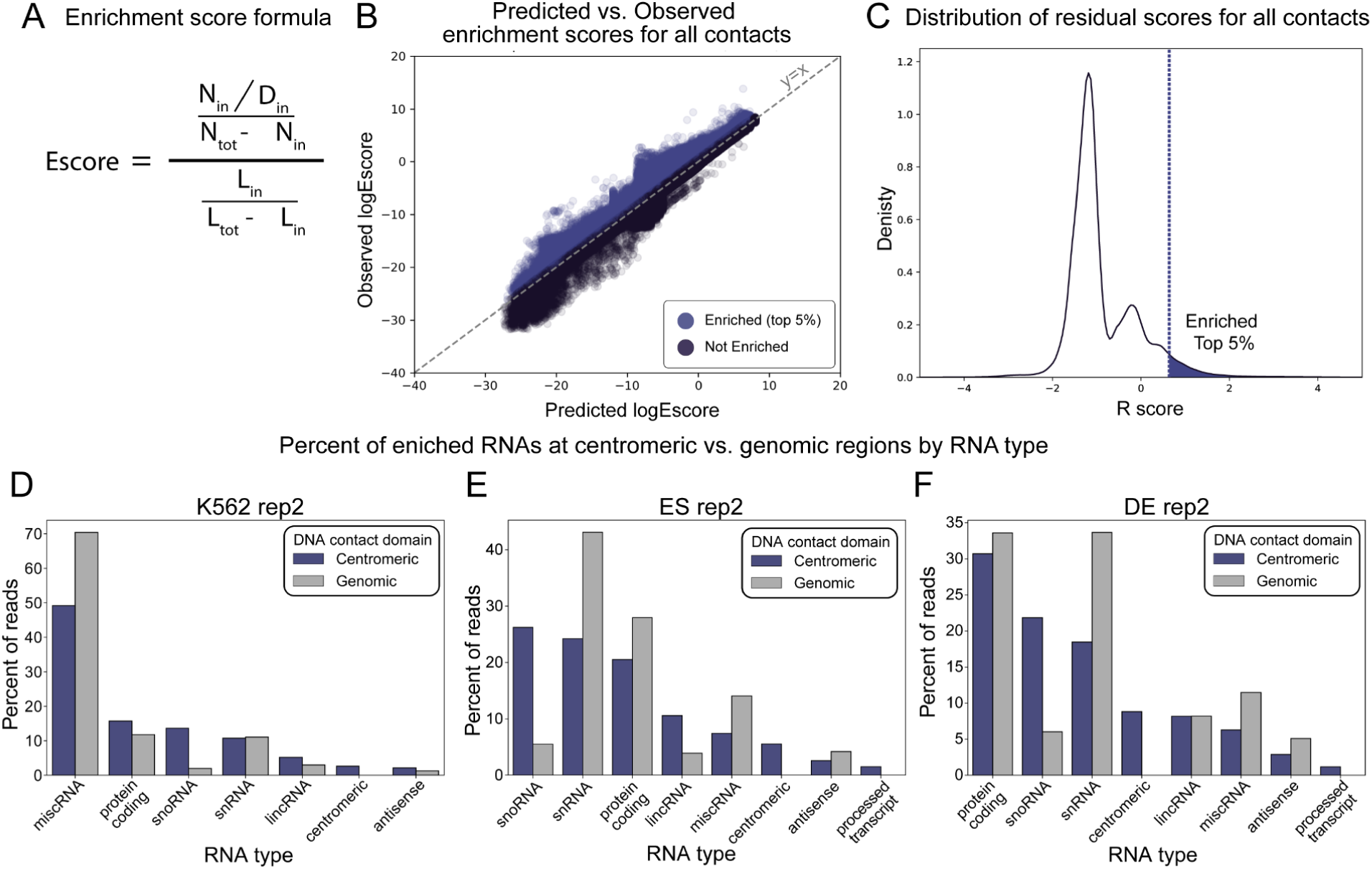
Enrichment score calculation and enriched RNA types at centromeres vs. genomic regions. **A.** Enrichment score calculation for each RNA-DNA contact. N_in_ represents the number of reads where a given RNA was found contacting a given DNA domain. N_tot_ is the total number of times that RNA was found in the entire dataset. D_in_ is the total number of DpnII sites in the DNA domain. L_i_n is the length of the DNA contact domain and L_tot_ is the total length of the genome. **B.** Scatter plot depicting observed (y-axis) and predicted (x-axis) enrichment scores (E scores) for all RNA-DNA contacts from K562 rep1. Mitochondrial and ribosomal RNA contacts were excluded. Residual scores were calculated as the difference between predicted and observed E scores and the top 5% were considered significantly enriched (purple) **C.** Distribution of residual scores. Top 5% (purple section denoted by vertical dotted line) defined as significantly enriched. **D-F.** Proportion of RNA reads enriched at centromeric (purple) or non-centromeric (gray) domains belonging to each RNA type in **D**. K562 replicate 2, **E.** ES replicate 2, and **F.** DE replicate 2.

**Supplemental Figure S5:**
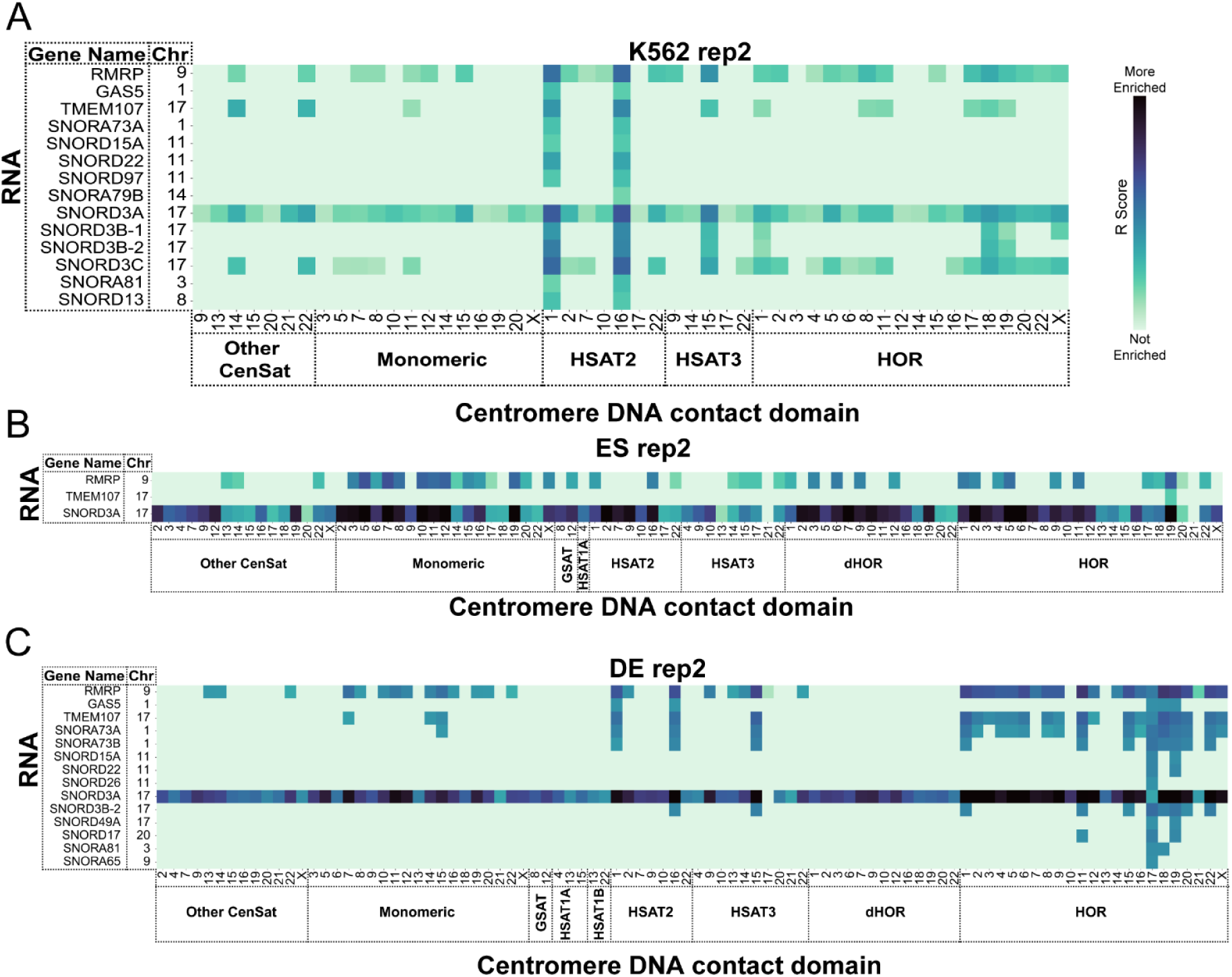
Nucleolar localized RNAs enriched at centromere DNA domains. Heatmap of R scores for nucleolar localized RNAs (x-axis) enriched at centromeric repeat domains (y axis) in **A.** K562 replicate 2, **B.** ES replicate 2, and **C.** DE replicate 2. DNA side reads were classified into centromeric repeat domains using CASK and are denoted on x-axis with domain type label and chromosome. RNA reads were classified by genomic alignment and denoted with gene name and the chromosome they originate from. Darker color indicates contacts with higher R scores (larger difference between observed number of reads and predicted) indicating stronger enrichment above background contacts. Mitochondrial RNAs, ribosomal RNAs, ribosomal DNA, reads that could not be uniquely assigned to a single repeat type, and all contacts with fewer than 0.2CPM were excluded from heatmap.

**Supplemental Figure S6:**
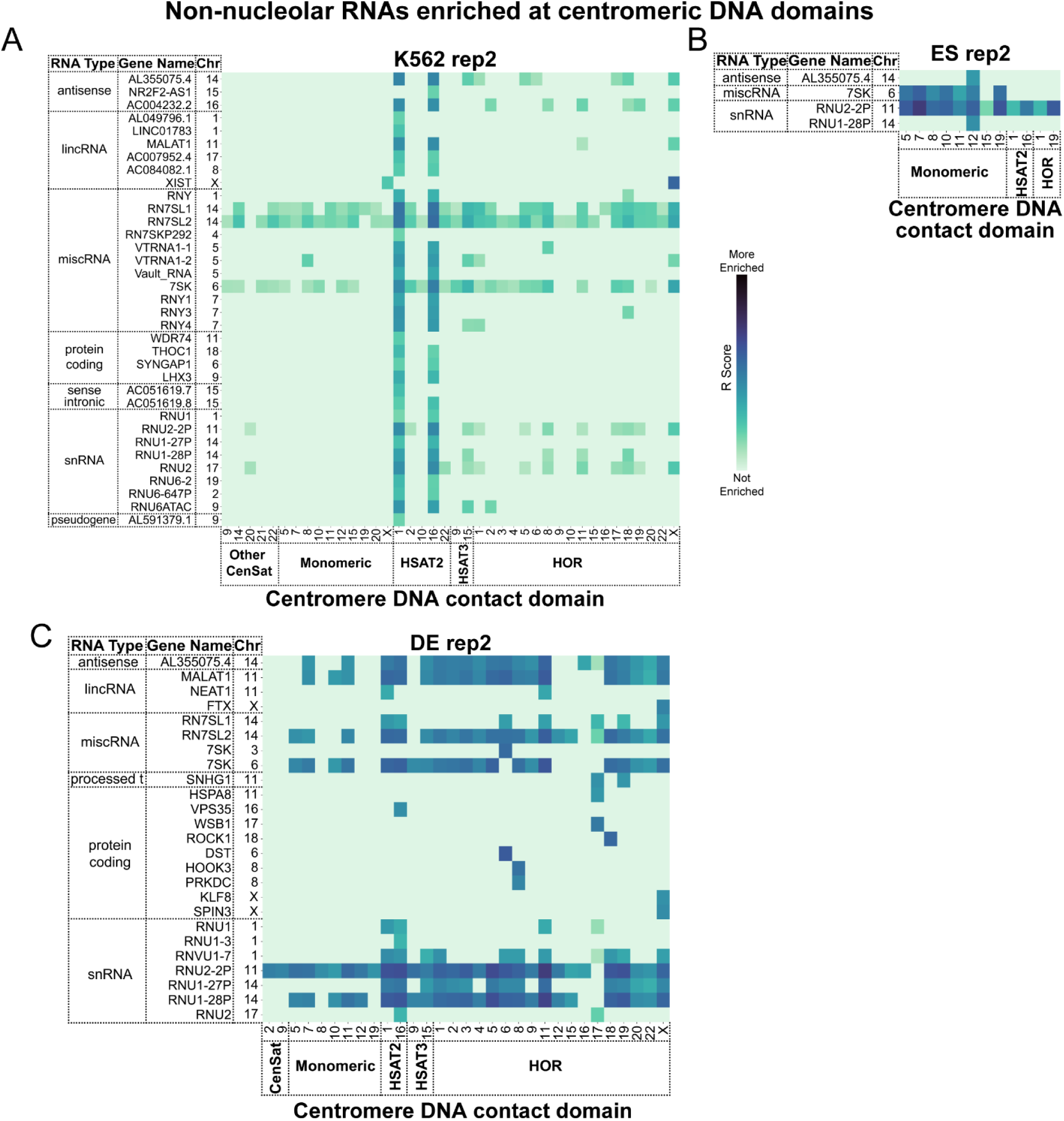
Non-nucleolar localized RNAs enriched at centromere DNA domains. Heatmap of R scores for all non-nucleolar localized RNAs (x-axis) enriched at centromeric repeat domains (y axis) in **A.** K562 replicate 2, **B.** ES replicate 2, and **C.** DE replicate 2. DNA side reads were classified into centromeric repeat domains using CASK and are denoted on x-axis with domain type label and chromosome. RNA reads were classified by genomic alignment and denoted with gene name and the chromosome they originate from. Darker color indicates contacts with higher R scores (larger difference between observed number of reads and predicted) indicating stronger enrichment above background contacts. Mitochondrial RNAs, ribosomal RNAs, ribosomal DNA, reads that could not be uniquely assigned to a single repeat type, and all contacts with fewer than 0.2CPM were excluded from heatmap.

**Supplemental Figure S7:**
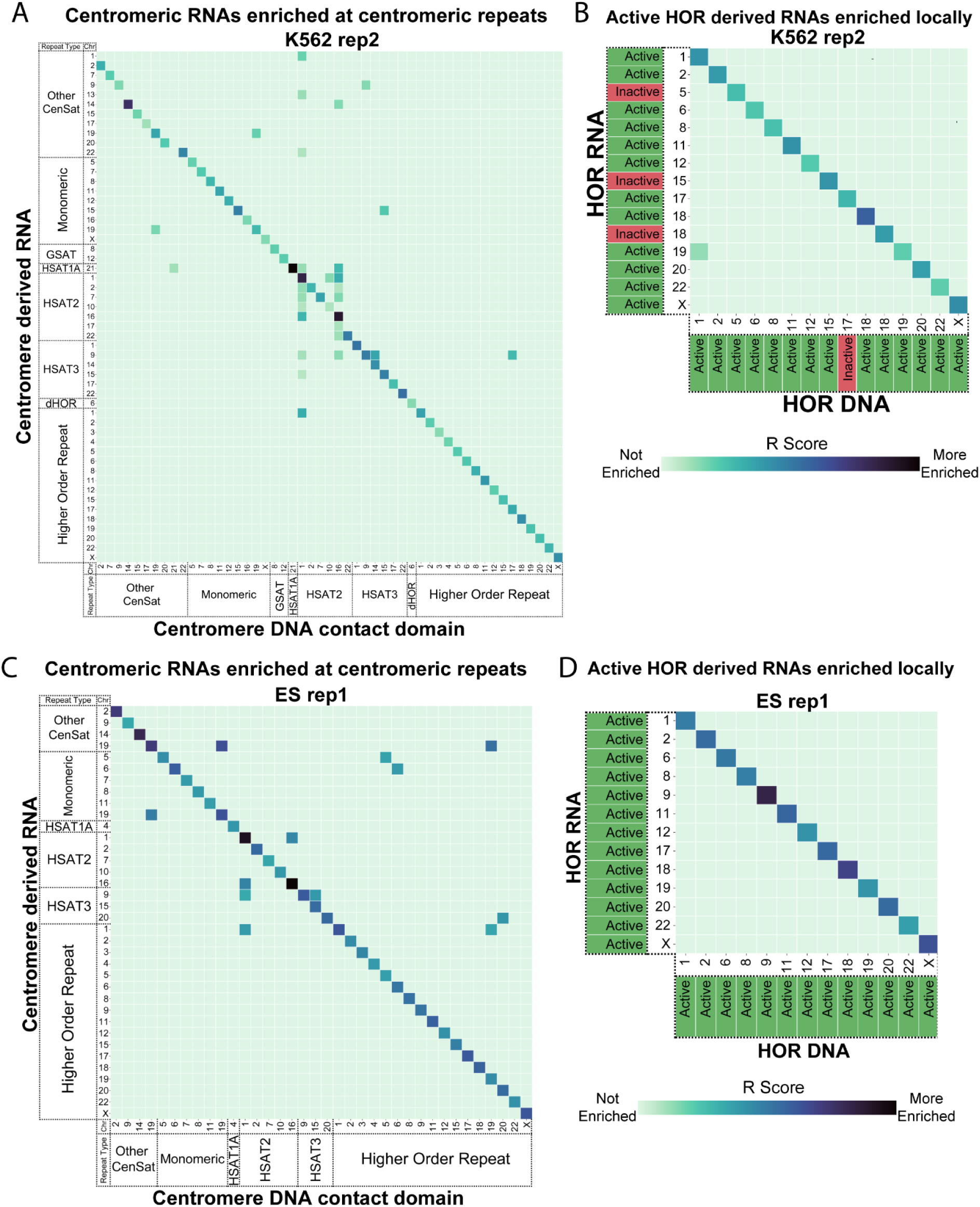
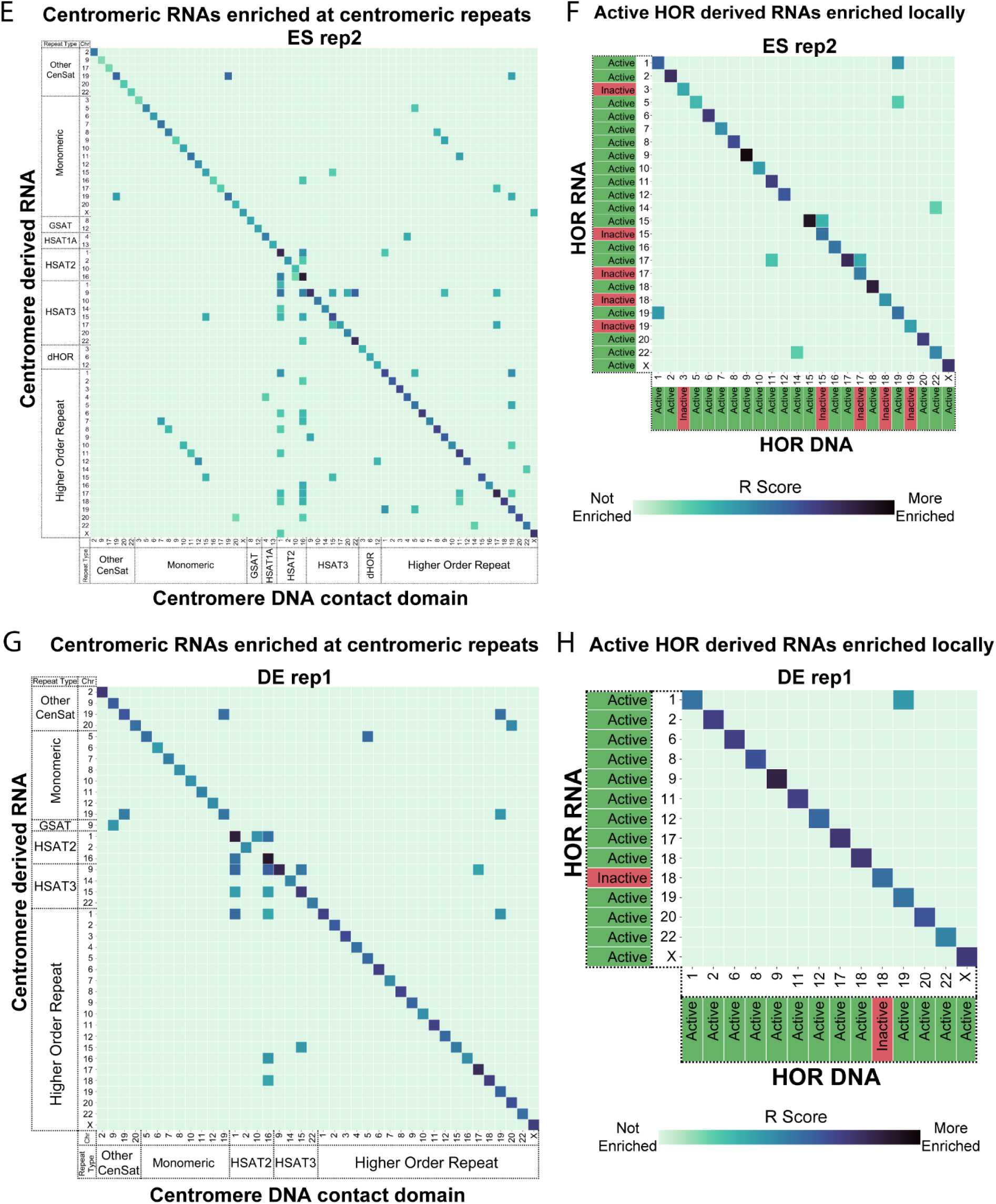
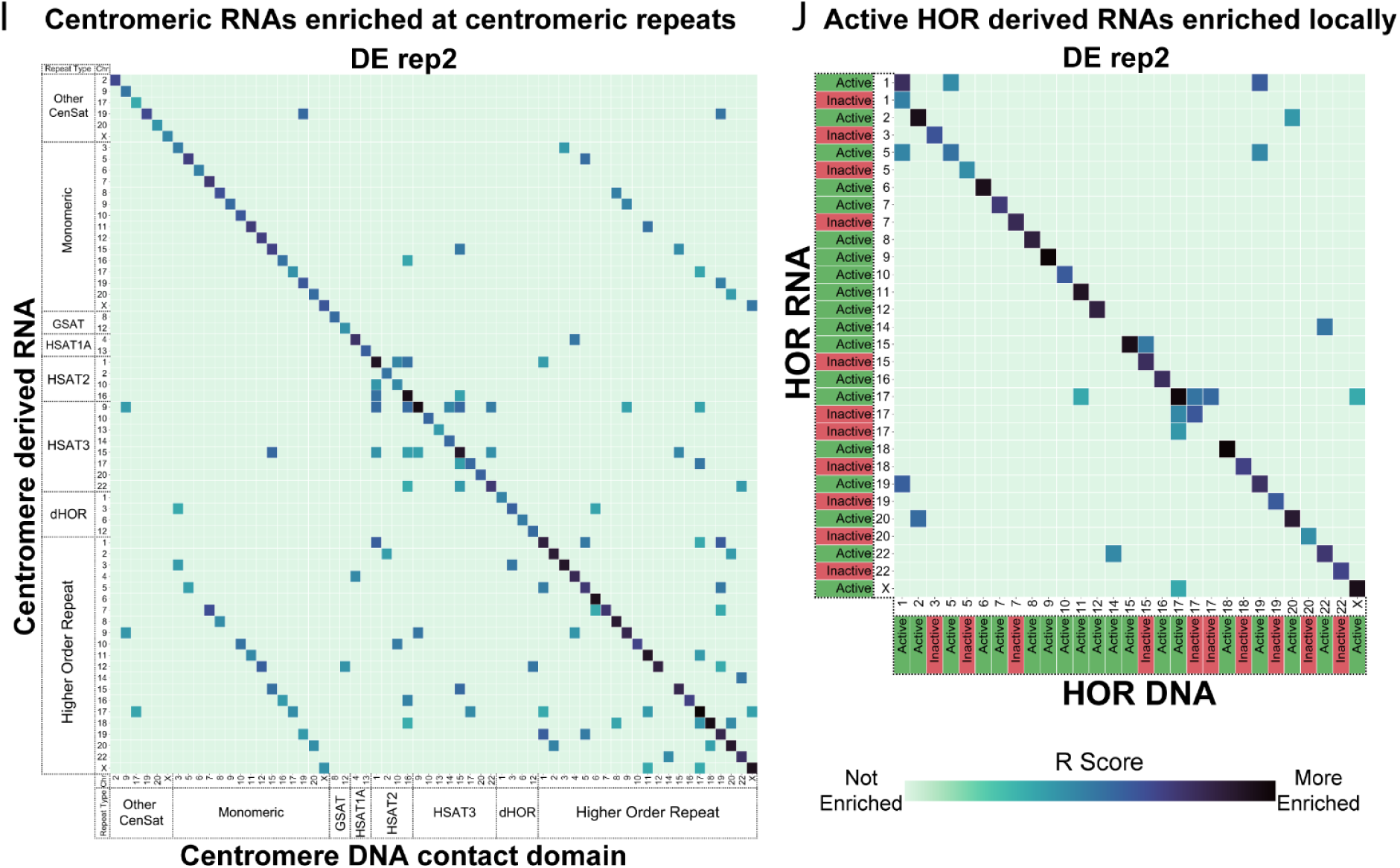
Centromere derived RNAs enriched at centromere DNA domains. As in Figure 4A,B, heatmaps of R scores for centromere derived RNAs (y-axis) enriched at centromere DNA domains (x-axis) in **A-B.**K562 replicate 2, **C-F** ES replicates 1 and 2, and **G-J.** DE replicates 1 and 2. DNA and RNA reads were classified using CASK and denoted with domain type label and chromosome. Darker color indicates contacts with higher R scores (larger difference between observed number of reads and predicted) indicating stronger enrichment above background contacts. Contacts with R scores below the 95th percentile were considered not significantly enriched and were excluded from heatmaps. Reads classified as ribosomal on either RNA or DNA side, reads that could not be uniquely assigned to a single repeat type, and all contacts with fewer than 0.1CPM of total reads were excluded from heatmap. **A,C,E,G,I.** Heatmap of R scores for class level assigned centromere derived RNAs enriched at centromere DNA domains in **A.** K562 replicate 2 **C.** ES replicate 1 **E**. ES replicate 2 **G.** DE replicate 1 **I**. DE replicate 2 **B,D,F,H,J**. Heatmap of R scores for repeat type level HOR derived RNAs enriched at HOR DNA domains in **B.** K562 replicate 2 **D.** ES replicate 1 **F**. ES replicate 2 **H.** DE replicate 1 **J**. DE replicate 2. Green label corresponds to T2T active HOR annotation denoting CENP-A containing HOR. Red, inactive label denotes HOR that does not contain CENP-A.

**Supplemental Figure S8:**
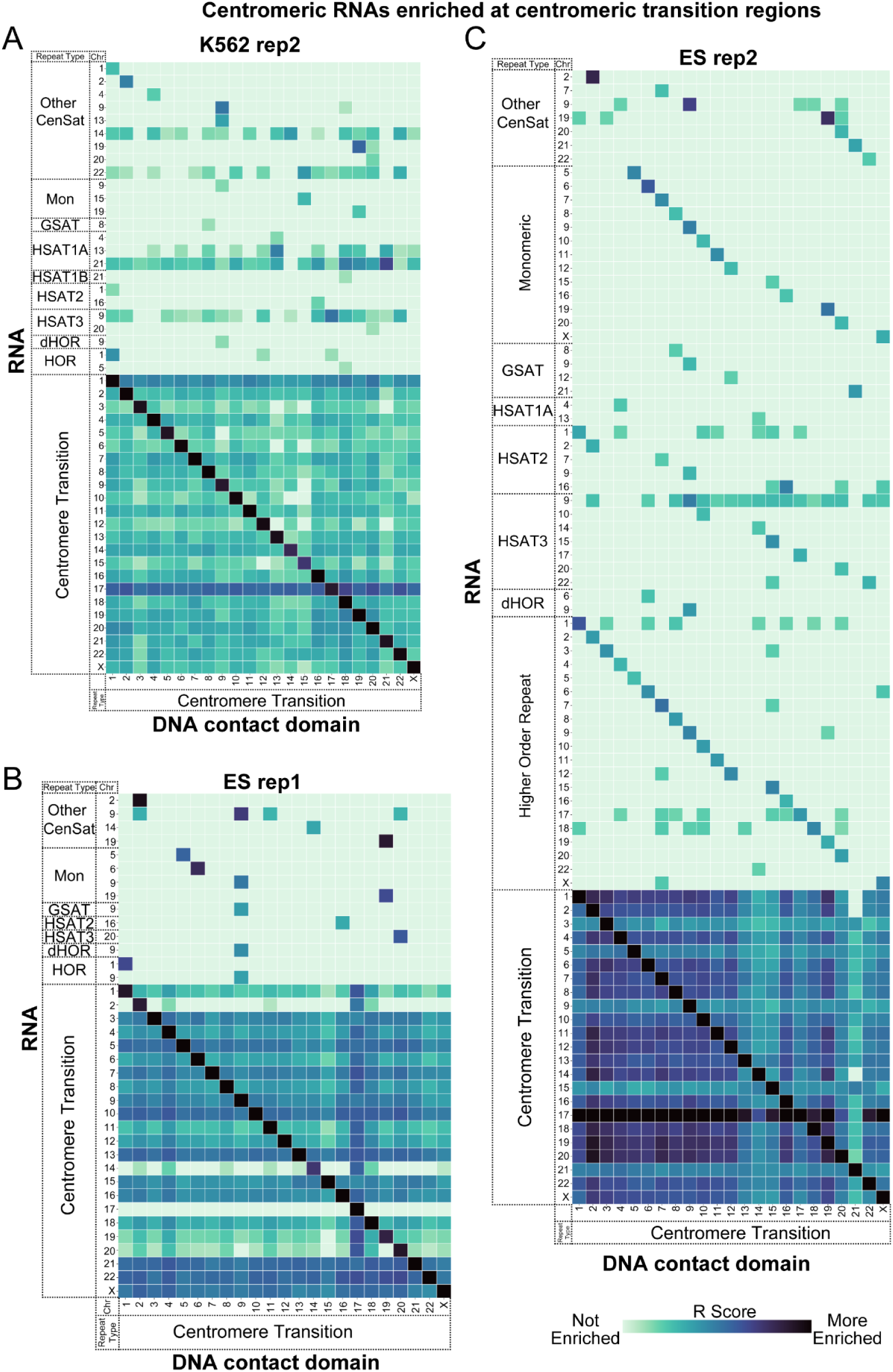
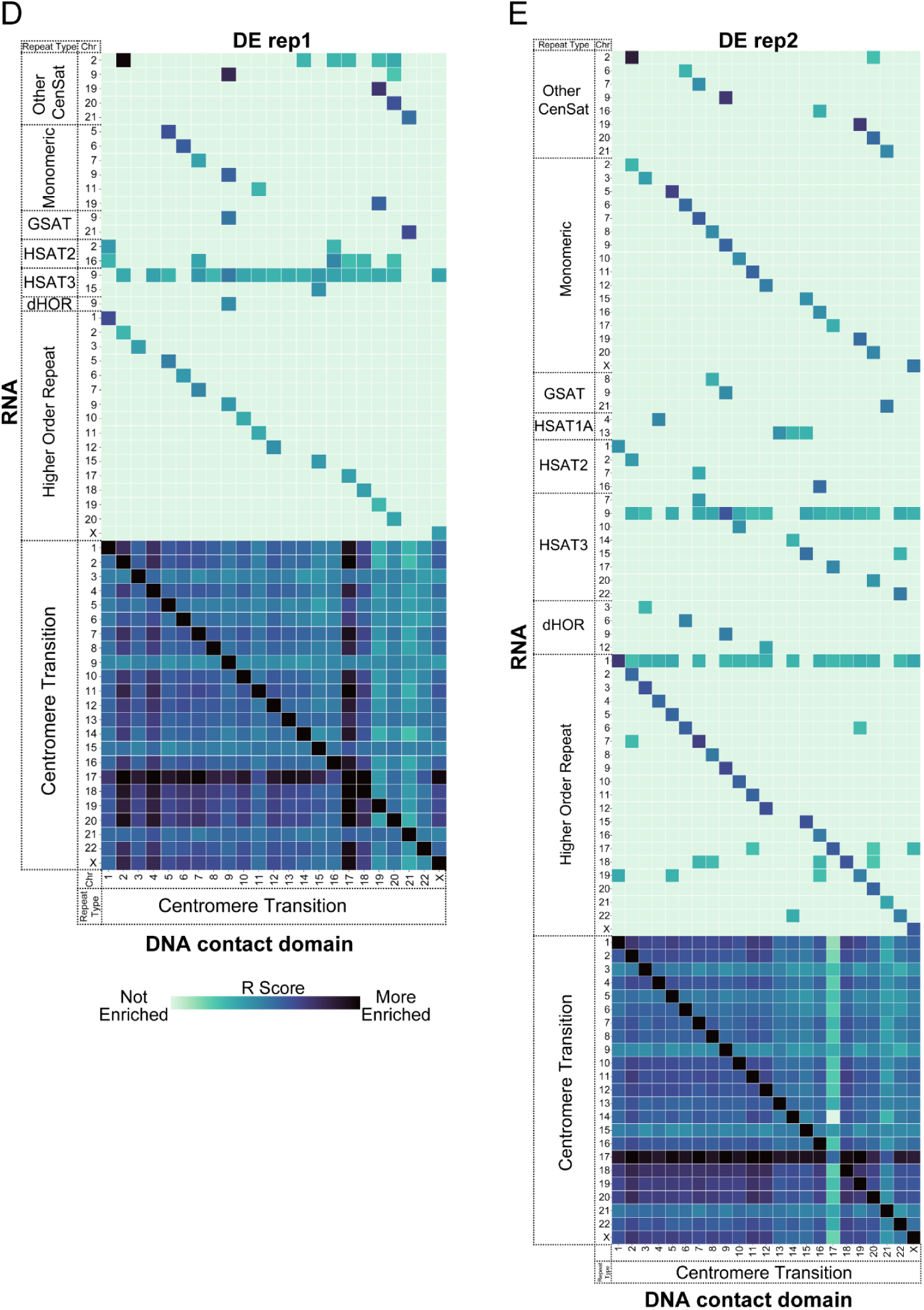
Centromere derived RNAs enriched at centromere transition DNA domains. As in Figure 4C: heatmaps of R scores for centromere derived and centromere transition (CT) derived RNAs (y-axis) enriched at CT DNA domains (x-axis) in **A.** K562 replicate 2 **B.** ES replicate 1 **C**. ES replicate 2 **D.** DE replicate 1 **E**. DE replicate 2. DNA and RNA reads were classified using CASK and denoted with domain type label and chromosome. Darker color indicates contacts with higher R scores (larger difference between observed number of reads and predicted) indicating stronger enrichment above background contacts. Contacts with R scores below the 95th percentile were considered not significantly enriched and were excluded from heatmaps. Reads classified as ribosomal on either RNA or DNA side, reads that could not be uniquely assigned to a single repeat type, and all contacts with fewer than 0.1CPM of total reads were excluded from heatmap.

**Supplemental Figure S9:**
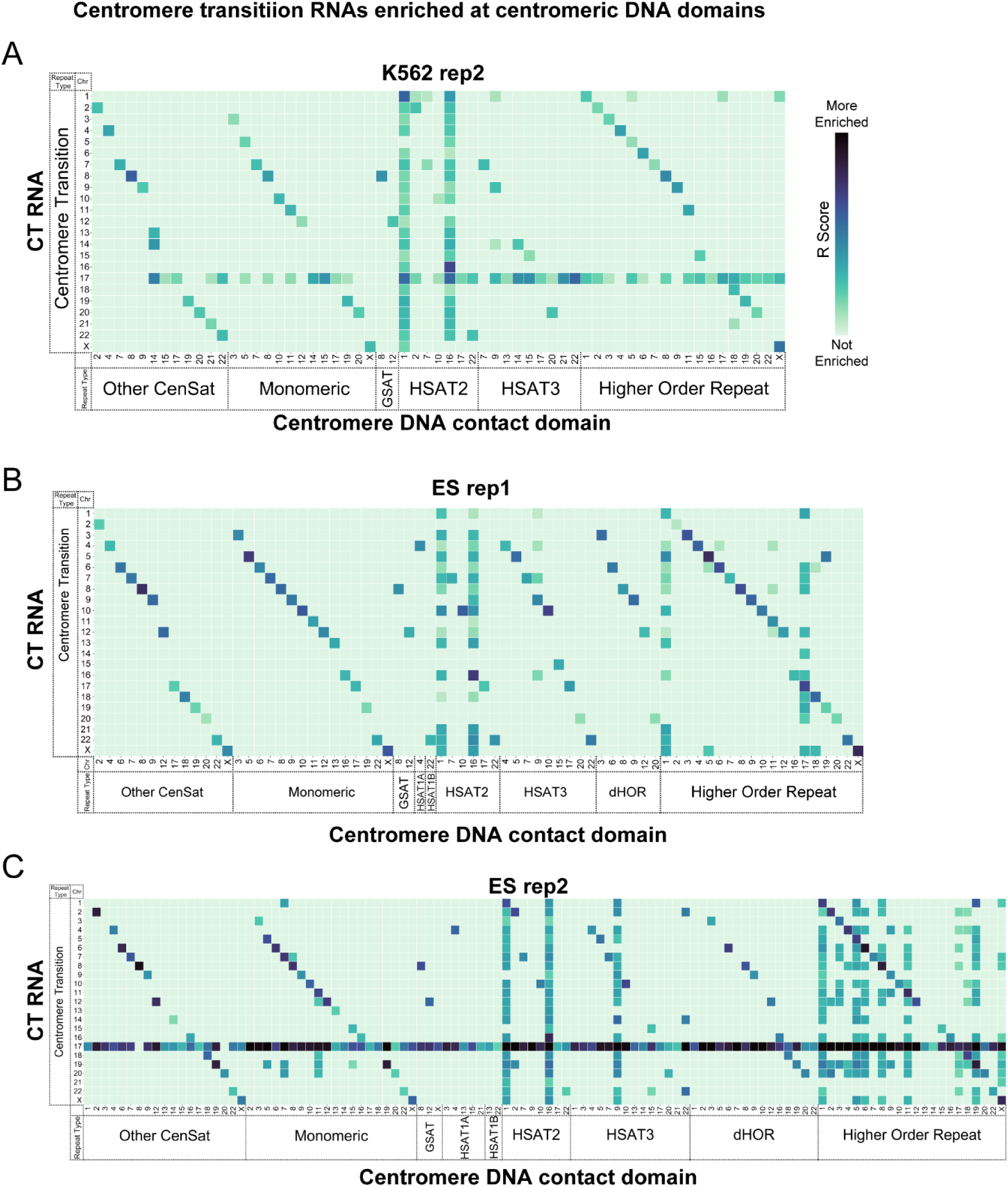
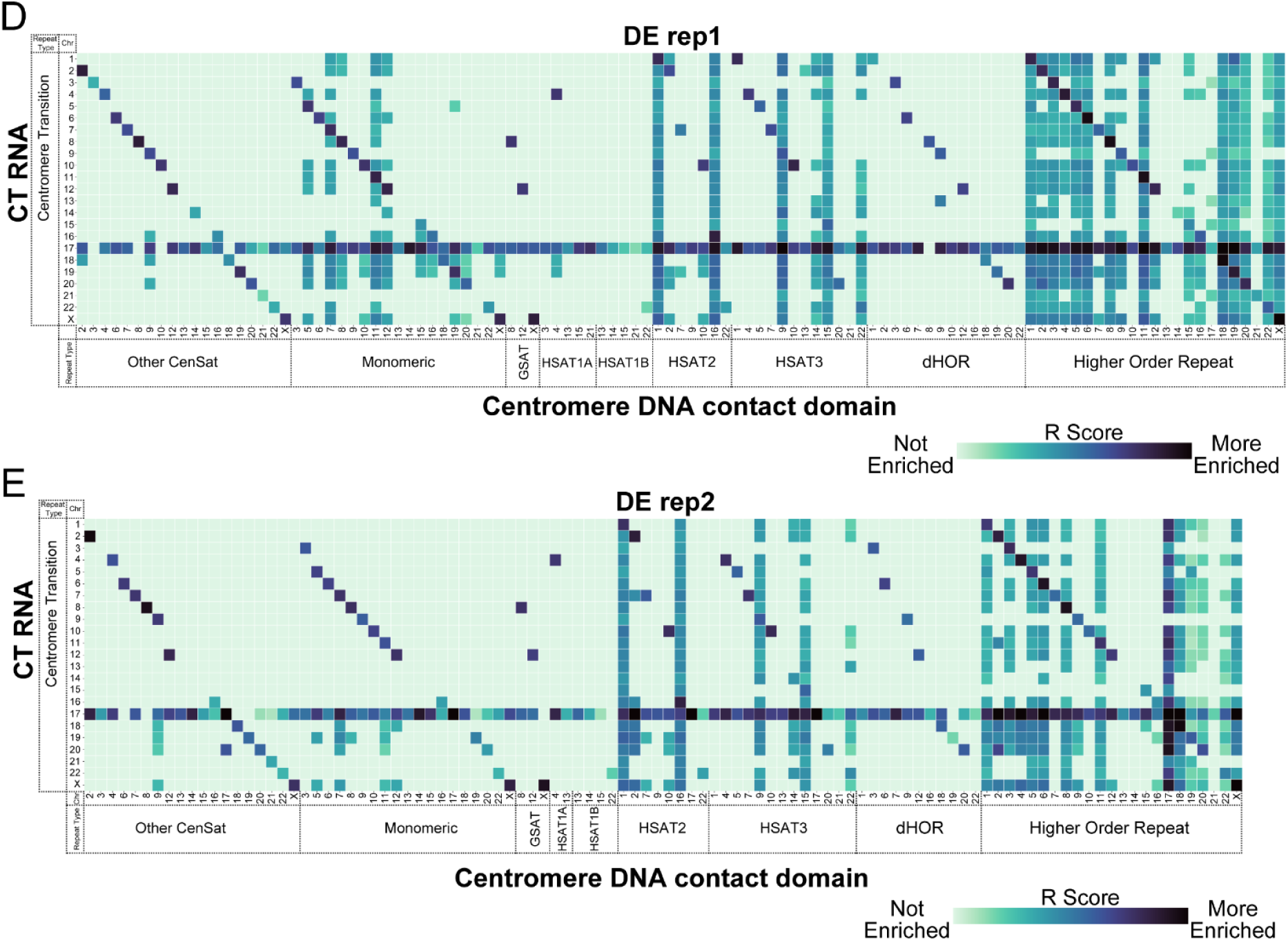
Centromere transition derived RNAs enriched at centromere DNA domains. As in Figure 4D, heatmaps of R scores for centromere transition region (CT) derived RNAs (y-axis) enriched at centromere DNA domains (x-axis) in **A.** K562 replicate 2 **B.** ES replicate 1 **C**. ES replicate 2 **D.** DE replicate 1 **E**. DE replicate 2. DNA and RNA reads were classified using CASK and denoted with domain type label and chromosome. Darker color indicates contacts with higher R scores (larger difference between observed number of reads and predicted) indicating stronger enrichment above background contacts. Contacts with R scores below the 95th percentile were considered not significantly enriched and were excluded from heatmaps. Reads classified as ribosomal on either RNA or DNA side, reads that could not be uniquely assigned to a single repeat type, and all contacts with fewer than 0.1CPM of total reads were excluded from heatmap.

## Notes

https://www.ncbi.nlm.nih.gov/geo/query/acc.cgi?acc=GSE298896

https://github.com/straightlab/

### Competing Interest Statement

The authors have declared no competing interest.

